# Circular RNAs in Lotus japonicus Responses to Nutrient Supply and Symbiotic Interactions

**DOI:** 10.1101/2025.08.28.672498

**Authors:** Delecia Utley, Asa Budnick, Simona Radutoiu, Heike Sederoff

**Author notes:** Corresponding author: Heike Sederoff.

## Abstract

Symbiotic relationships, such as those formed between legumes and rhizobia or arbuscular mycorrhizal (AM) fungi, function by enhancing the nutrient uptake into the plant. The establishment and coordination of the symbiotic interactions requires changes in gene expression in the host and microbe. Circular RNAs (circRNAs) can function through sponging of microRNAs (miRNAs), resulting in changes in transcript abundances. We identified 15,252 unique nuclear circRNAs in *Lotus japonicus* under different nutrient conditions and symbiotic interactions with rhizobia or AMF. Our results revealed treatment-specific circRNAs and circRNAs originating from key genes in the Common Symbiosis Pathway, suggesting their potential role in the establishment of these symbioses. We validated select circRNAs potentially involved in the regulation of symbiosis and predicted miRNA recognition elements (MREs) that were only created by the backsplice junction of circRNAs. Backsplice-generated MREs represent an additional mechanism through which circRNAs may modulate abundances and translation of mRNAs. Our sequencing approach using random hexamer primers also enabled us to simultaneously characterize the transcriptome of the symbionts and host.

## 1 Introduction

Harnessing the power of symbiotic relationships formed between plants and beneficial microbes will be key to advancing sustainable agriculture. These symbiotic relationships enhance nutrient uptake, improve soil health, and reduce the reliance on chemical inputs. Legumes are unique in their ability to form symbiotic relationships with both phosphorus-acquiring arbuscular mycorrhizae and nitrogen-fixing rhizobia. *Lotus japonicus* has been studied as a model legume to understand the evolution, signaling, and development of those symbiotic interactions.

Perception is the first step in the development of symbiotic relationships. Plants excrete secondary metabolites that are recognized by AM fungi and rhizobia, resulting in the production of lipochitooligosaccharide (LCOs) and Nod factors from AM and rhizobia, respectively. In *L. japonicus*, the Nod factor receptors, *NFR1* and *NFR5,* are responsible for recognizing these signals (Broghammer et al., 2012; Murakami et al., 2018).

Downstream of recognition, the symbiotic relationships in legumes with AM or rhizobia are orchestrated by a shared molecular pathway in plants, known as the Common Symbiosis Pathway (CSP), which includes several genes that are involved in nodulation and colonization of AM fungi. CSP genes that are shared among the two symbiotic relationships include the *Symbiosis Receptor Like Kinase* (*SymRK)* (Stracke et al., 2002)*, CASTOR* and *POLLUX* (Charpentier et al., 2008)*, Calcium and Calmodulin-dependent protein Kinase (CCaMK)* (Gleason et al., 2006), and *CYCLOPS* (S. Singh et al., 2014; Yano et al., 2008).

While the genetic framework governing these processes is well-established, emerging research suggests that RNA, particularly non-coding RNA, plays a crucial role in regulating the expression of some symbiosis-related genes. Non-coding RNAs, such as microRNAs (miRNAs), transfer RNA (tRNA) derived small RNA fragments (tRFs), and circular RNAs (circRNAs), are increasingly recognized as important regulators of gene expression, including the fine-tuning of plant responses to microbial signaling (Okuma & Kawaguchi, 2021; B. Ren et al., 2019; Tsikou et al., 2018). Understanding the role of non-coding RNA in these pathways offers new insights into how plants manage complex symbiotic relationships post-transcriptionally. For example, miR2111 is involved in the systemic regulation of nodulation in *L. japonicus* (Tsikou et al., 2018). Two other miRNAs, miR397 and miR171, are systemically regulated and target a transcription factor involved in organogenesis of nodules in *L. japonicus* and AM colonization in Medicago (De Luis et al., 2012). Rhizobial tRFs were found to regulate host genes involved in nodulation by affecting the RNA interference machinery, a novel mechanism showing regulation of host genes by the symbiont (B. Ren et al., 2019). More recently, circRNAs have emerged as crucial players in plant responses to abiotic and biotic conditions (Lv et al., 2020; P. Ma et al., 2021; Wu et al., 2020; Xia et al., 2023; Z. Zeng et al., 2023).

CircRNAs are single-stranded, covalently closed RNA molecules, distinguishable only by their back-splice junction (BSJ) (Zhang et al., 2024). The BSJ is formed by the ligation of the 3’ and 5’ ends of linear RNA, making these circular molecules inaccessible to exonucleases. In plants, there are several studies focusing on the discovery of circRNAs under different abiotic and biotic conditions (Lv et al., 2020; P. Ma et al., 2021; Wu et al., 2020; Z. Zeng et al., 2023). CircRNAs have been identified in response to AM fungi in tomato (Z. Zeng et al., 2023), to nodulation in *Phaseolus vulgaris* (Wu et al., 2020), and to nitrogen fertilization or availability in bamboo, maize, wheat, and poplar (Liu et al., 2020; P. Ma et al., 2021; Y. Ren et al., 2018; Zhu et al., 2023). CircRNAs were detected in soybean experiencing low phosphorus conditions (Lv et al., 2020). Some studies identify circRNAs involved in plant defense and immunity responses (Fan et al., 2020; Xia et al., 2023). Collectively, these findings provide key insights into the potential role of circRNAs in plant symbiosis and nutrient signaling.

One well-documented function of circRNAs in both animals and plants is transcriptional regulation. In animals, some circRNAs enhance their own gene expression by interacting with the transcriptional machinery (Guria et al., 2020), while others bind to their cognate mRNA to suppress expression (Guria et al., 2020; Li et al., 2017). In plants, a circRNA from exon 6 of the *SEPALLATA3* (*SEP3*) gene in *Arabidopsis thaliana* forms an R-loop on the parent gene; transgenic overexpression of the circRNA leads to an increased abundance of the exon-6-skipped alternative splice (AS) variant and a floral phenotype (Conn et al., 2017).

CircRNAs are well known for their role as miRNA sponges. A classic example in animals is ciRS-7, which binds to miR-7, a key regulator in various cancers (Guria et al., 2020; Kefas et al., 2008). CiRS-7 contains multiple binding sites for miR-7, many of which are conserved across vertebrates, effectively sequestering miR-7 and preventing its interaction with mRNA targets (Guria et al., 2020; Zhang et al., 2024). In plants, circR5g05160 enhances *Magnaporthe oryzae* blast resistance in rice through miRNA sponging (Fan et al., 2020).

CircRNA’s ability to compete with linear mRNA for miRNA binding forms a competing endogenous RNA (ceRNA) network that coordinates expression within a pathway or between networks. The ceRNA model suggests that circular RNAs containing shared miRNA response elements (MREs) can compete for miRNA binding, thereby regulating each other’s expression levels (Salmena et al., 2011). This hypothesized mechanism creates a miRNA-mediated regulatory network linking different RNA species, including mRNAs, long non-coding RNAs (lncRNAs), and circRNAs (Salmena et al., 2011). By sequestering miRNAs, circRNAs can protect target mRNAs from degradation or translational inhibition. Several plant studies have predicted that these ceRNA networks function under different conditions (Wang et al., 2022; Xu et al., 2023; Zhou et al., 2022). In common bean, this network was predicted to regulate gene expression during nodule formation and nitrogen fixation. During nitrogen-fixation, more miRNAs, circRNAs, and mRNAs were predicted to be interacting or in competition with each other compared to during nodule formation (Wu et al., 2020). During AM symbiosis in tomato, a predicted ceRNA network showed 10 differentially expressed miRNAs could target 331 RNAs, the majority being mRNAs, followed by lncRNAs and circRNAs (Z. Zeng et al., 2023). Understanding ceRNA interactions across different biological systems provides insight into the complexity of gene expression regulation.

Our study identified the circRNA landscape of *L. japonicus* under nutrient stress and symbiotic interactions. A key novelty of our work is the identification of circRNAs acting in the CSP pathway during rhizobia and AMF symbiosis, as well as a broader set of circRNAs associated with nutrient signaling. We developed a novel approach to map miRNA to potential new miRNA recognition elements (MREs) generated by the circRNA BSJ, constructing a competitive endogenous RNA (ceRNA) network that sheds light on the regulatory roles of circRNAs in plant responses to symbiotic interactions. By analyzing differentially expressed genes (DEGs) and employing multiple circRNA detection methods, we further demonstrated the impact of sequencing depth and symbiotic partners on the circRNA landscape. Lastly, we validated key circRNAs, including those from *CCaMK*, and observed tissue-specific expression patterns. Our deep sequencing and use of random primers for cDNA synthesis provided simultaneous transcriptomes from lotus and rhizobia, which allows a rare snapshot of coordinated responses between both organisms.

## 2 Materials and Methods

### 2.1 Plant and Inoculant Material

*Lotus japonicus* (ecotype Gifu B-129) seeds were scarified, surface-sterilized in 7% bleach (10 min), rinsed five times in sterile water, and imbibed overnight at 4 °C in the dark. Seeds were stratified on moist sterile filter paper for 3 days at 22–23 °C in the dark, then transferred to a 16 h light/8 h dark cycle (50–120 µmol m⁻² s⁻¹) for 7 days.

Seedlings were grown on vertical plates with ½ MS (high-nutrient; Hn) or custom low-nutrient (Ln) MS media (Phytotech Labs) containing 1.4% agar. The Hn media contained 30mM total nitrogen (N) and 0.625 mM phosphate (PO₄³⁻), while the Ln media contained 2 mM N and 0.0625 mM PO₄³⁻ (see Supplemental File 1 for full recipe). A 30µm piece of nylon mesh was placed over the agar, and stainless-steel zip ties covered in black tape with punched holes for seedlings were positioned to block light while holding 10 seedlings per plate. Roots were positioned on the nylon mesh in contact with the media. Plates were sealed with parafilm and micropore tape, wrapped in black sleeves, and incubated vertically in the growth chamber.

After one week, seedlings on Ln media were inoculated with either *Mesorhizobium loti* (500 µL; OD₆₀₀ = 0.02 in Ln media) or ∼100 pre-germinated *Rhizophagus irregularis* spores (Premier Tech) in 500 µL sterile water. AMF spores were pre-germinated for 7 days at 26 °C as described previously (Fernández et al., 2019; Hornstein et al., 2024). *M. loti was* cultured in YMB medium for 3 days at 30°C with shaking (100 rpm). Control plates received sterile media. After inoculation, plates rested flat for 1hr before being returned to growth conditions for 3 weeks. Plants were grown in four independent experimental batches. In total, 48 plates were used, with approximately equal numbers of replicates from each batch to ensure balanced representation.

Before harvest, plates were imaged for phenotyping. Roots were excised with sterile razor blades, and tissues from five plants from the same plate were pooled per tube with Zirconia/Silica and metal beads, flash frozen in liquid nitrogen, and stored at –80 °C.

### 2.2 Phenotyping

Root lengths were measured in FIJI (ImageJ) by tracing the primary root using the segmented line tool and setting the scale based on a 20 × 20 cm grid in the image. Lateral and tertiary roots were counted manually to assess root branching. Shoot height was estimated from the same grid in 0.5 cm increments. For Rhizobia-treated plants, pink nodules were counted per plant, and averages were calculated per biological replicate. Data were analyzed in R using ANOVA followed by Tukey’s HSD post hoc test when p < 0.05.

### 2.3 RNA Extraction and Sequencing

Root tissue from five pooled plants was homogenized twice at 30 Hz for 45 sec using a tissue lyser. Total RNA was extracted using the Purelink RNA Mini Kit (Invitrogen). Each biological replicate consisted of pooled RNA from 10 plants (two tubes, combined through a single column). RNA was DNase-treated (TURBO DNase, Invitrogen), quantified via Qubit fluorometry, and assessed for quality with the Bioanalyzer 2100; only samples with RIN ≥8 were sequenced.

Ribosomal RNA was depleted using the QIAseq FastSelect–rRNA Plant Kit (Qiagen). Libraries were prepared with the NEBNext® Ultra™ II Directional RNA Library Prep Kit and multiplexed with NEBNext® Dual Index Primers (NEB). Sequencing was performed at NCSU Genomic Sciences Lab on an Illumina NovaSeq S4 platform (150bp paired end, 40-100 million reads/sample across two runs). Read quality was assessed with FastQC (Andrews, 2010). Trimmomatic (Bolger et al., 2014) was used to remove adapters and low-quality reads (sliding window: 4 bp, phred ≥20, min length 30 bp); unpaired reads were discarded. Resulting fastq files were used for circRNA and linear RNA analysis.

### 2.4 RNA-seq alignment and gene expression

Trimmed paired reads were aligned using BBSplit (Bushnell, 2014) against three reference genomes: *Lotus japonicus* Gifu v1.2 assembly (Genbank: GCA_012489685.2), *Mesorhizobium loti* (Genbank: GCA_012913625.1), and *Rhizophagus irregularis* (Genbank: GCA_000439145.3). High stringency settings (minimum identity = 0.97, ambiguous reads removed) ensured accurate organism-specific read assignments. Ribosomal RNA counts were used to verify symbiont presence in each sample. Reads assigned to each genome were quantified at the gene level using featureCounts (Liao et al., 2014) with default settings. Counts for *M. loti* 5S, 16S, and 23S ribosomal RNA and *R. irregularis* 18S, 40S, and 60S rRNA genes were summed to confirm sample treatment identities.

Reads aligned to the *L. japonicus* genome were individually counted at the gene level using featureCounts with default settings. Counts for each sample were merged into a single count matrix. Genes with zero counts or low abundance across all samples (<10 total reads) were filtered out. Differential Gene Expression (DGE) analysis was performed with edgeR (v4.2.1) (Robinson et al., 2010). Counts were normalized, and a group matrix without an intercept was used to allow more flexible comparisons between treatments (i.e., no pre-defined control group). A generalized linear model (GLM) was applied, and edgeR’s automated dispersion estimation was used to determine the optimal transformation of normalized counts. Likelihood ratio tests were performed for relevant pairwise contrasts. Contrasts followed a “treatment vs control” format, where positive log fold changes indicate upregulation in the treatment relative to the control. All treatments were compared using single pairwise contrasts.

### 2.5 GO term Analysis

Gene ontology (GO) enrichment analysis for Biological Process terms was performed using the topGO (Adrian Alexa, 2017) R package with the *L. japonicus* Gifu v1.2 GO annotations from Lotus Base. Enrichment was tested using the ‘weight01’ algorithm and Fisher’s exact test via the runTest function. The background gene universe included 13,654 annotated genes. Terms with *P* < 0.001 were considered significant, and the top 50 were reported.

### 2.6 CircRNA Analysis

CircRNAs were identified using two independent pipelines: CIRI2 (v2.1.1) (Gao et al., 2018) and CLEAR (v1.0.1) (X.-K. Ma et al., 2019). Trimmed reads were aligned to the *L. japonicus* Gifu v1.2 genome using BWA-MEM (v0.7.17) with default parameters and a t-19 alignment score cutoff for CIRI2. For CLEAR, reads were aligned with HISAT2 (v2.2.1) and analyzed using default settings to detect circRNAs and calculate circRNA:linear RNA ratios. Linear and circular reads were normalized by read depth and transcript length and quantified in fragments per billion mapped bases (FPB). The CIRCscore was computed as the FPB ratio of circRNA to its corresponding linear transcript. Outputs from both pipelines were processed with custom R scripts for filtering, clustering, and summarization.

### 2.7 ceRNA networks

Raw fastq files were processed with CIRI-Full (v2.0) pipeline (Zheng & Zhao, 2018) to assemble full-length circRNA sequences. To test for back-splice generated miRNA recognition elements (bsgMREs), circRNA sequences were represented in the query both as the original full sequence from CIRI-full and as an offset version of that sequence made by repositioning the last 40 nt to the sequence start. Both original and ‘offset’ circRNA sequences were analyzed for predicted miRNA recognition elements (MREs) using psRNATargetV2 (Dai & Zhao, 2011) with *L. japonicus* miRNAs from miRBase (Kozomara & Griffiths-Jones, 2014). In cases where the ‘offset’ circRNA sequences were predicted to have distinct MREs compared to the original sequence, these new MREs were termed bsgMREs. For linear transcripts, we used StringTie (-- merge) (Pertea et al., 2015) to collapse transcripts across all samples into a unified reference set, which were then analyzed for miRNA response elements using psRNATargetV2 under the same parameters.

### 2.8 RT-PCR Validation and Sanger Sequencing

Total RNA was extracted from 50–100 mg of ground *L. japonicus* root tissue using a modified CTAB protocol (Chang et al., 1993) followed by TRIzol purification. Briefly, pre-warmed CTAB extraction buffer with 5% β-mercaptoethanol was added to the tissue, incubated at 65 °C, and centrifuged at 15,000 × *g* for 10 min. The aqueous phase underwent two chloroform (24:1) extractions before isopropanol precipitation at 4 °C for 1 hr. The RNA pellet was washed twice with 70% ethanol, resuspended in TRIzol, and re-extracted with chloroform, followed by a second isopropanol precipitation. The final pellet was washed, dried, and resuspended in 37 °C nuclease-free water. Each sample represented RNA pooled from 15 plants.

cDNA synthesis was performed using DNase-treated total RNA and SuperScript IV Reverse Transcriptase with random hexamers (Thermo Fisher Scientific), following the manufacturer’s instructions. Divergent primers were designed to span the back-splice junction (BSJ) and maximize internal circRNA sequence coverage. PCR amplification was performed using OneTaq Hot Start 2X Master Mix (New England Biolabs) on cDNA (10% of total volume) and on 100 ng of RNase A-treated genomic DNA (gDNA) as a control for false positives. PCR cycling followed manufacturer’s guidelines, with a 30-second extension for circRNAs. Amplicons were visualized via agarose gel electrophoresis. PCR products of expected size were gel-excised, purified, and submitted for Sanger sequencing to confirm the BSJ. Primer sequences are listed in Supplemental File 2.

All custom analysis scripts are available at: https://github.com/SederoffLab/Lotus_n_Friends

All raw sequencing data generated in this study are available through the NCBI Sequence Read Archive (SRA) under accession number PRJNA1220156.

## 3 Results

### 3.1 Experimental Design and Phenotypic Response

*Lotus japonicus* was grown under four conditions: high nutrient (Hn) (30mM total N and 0.625 mM PO₄³⁻), low nutrient (Ln) (2 mM total N and 0.0625 mM PO₄³⁻), Ln with *Mesorhizobium loti* inoculation (Rhiz), and Ln with arbuscular mycorrhizal fungi *Rhizophagus irregularis* (AM) inoculation. After a three-week interaction, plants showed distinct phenotypes (Figure 1A-B). Root length was significantly greater in Hn compared to Rhiz (p<0.05; Supplemental Figure 1A-B), while lateral root number did not differ across treatments (Supplemental Figure 1C). Shoot height varies significantly with Hn and Rhiz differing from each other and from Ln and AM, which were similar (Supplemental Figure 1D-E). Rhizobia inoculated plants formed an average of three nodules (Supplemental Figure 1F).

**Figure 1.**
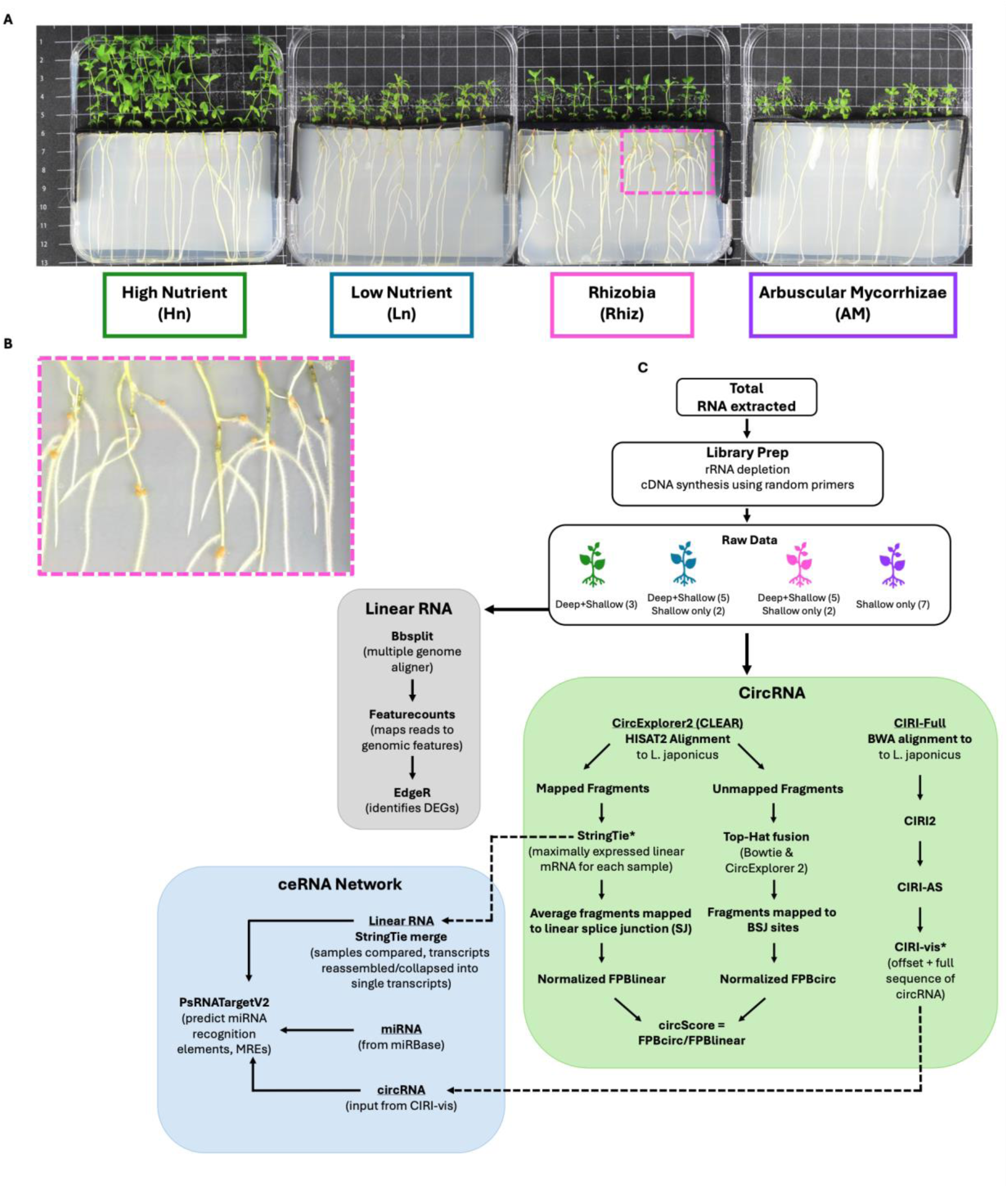
Overview of experimental design. (A) Representative *Lotus japonicus* plates from each treatment: high nutrient (Hn), low nutrient (Ln), Rhizobia (Rhiz), and arbuscular mycorrhiza (AM). Plants were grown for 38 days, including 21 days of interaction with *M. loti* (Rhizobia), *R. irregularis* (AM), or mock inoculation (Hn, Ln). (B) Close-up of an active nodule from a Rhizobia-treated plant. (C) Experimental schematic: rRNA-depleted libraries were sequenced at shallow (40M reads/sample) and deep (100M reads/sample) depths. Replicate numbers varied by depth. High-quality (FASTQC > 30), trimmed reads from both sets were combined for analysis.

Total RNA was extracted from biological replicates composed of 10 plants per treatment. Ribosomal RNA-depleted libraries were prepared using random primers to capture linear and circular RNAs. Initial “shallow” sequencing targeted 40 million 150bp paired end reads per sample and included 3 high nutrient (Hn) and 7 low nutrient (Ln), rhizobia (Rhiz), and arbuscular mycorrhizal (AM) replicates. Because circular RNAs are low in abundance, we hypothesized that greater depth would improve detection. Using the same libraries, we performed “deep” sequencing targeting 100 million reads per sample for 3 Hn, 5 Ln, and 5 Rhiz replicates. (Figure 1C).

Following filtering and adapter trimming, samples from shallow and deep sequencing were combined, resulting in 3 Hn, 7 Ln (2 shallow only), 7 Rhiz (2 shallow only), and 7 AM (shallow only) replicates. Sequencing depths ranged from 44M to 290M reads, averaging 259M for Hn, 211M for Ln, 203M for Rhiz, and 52M for AM (Supplementary Figure 2A, B; Supplemental File 3). BBsplit (Bushnell, 2014) was used to confirm symbiont presence (Supplemental Figure 2C-E; Supplemental File 3). Results confirmed replicate identities and showed lower AM read counts compared to Rhiz, likely due to the shallow sequencing depth of the AM treatment in this study, which may have limited detection of AM colonization at the 3-week timepoint (Xue et al., 2015). Despite sequencing depth and incomplete colonization, some AM-specific molecular responses were detected. Deep sequencing also enabled detection of the *M. loti* transcriptome, providing insight into symbiont gene expression, though comparisons are limited due to the absence of a host-free control (See section 3.3).

### 3.2 Low Nutrient and AM Treatments are Similar in Transcriptomic Response

We first assessed how treatment conditions impact the *L. japonicus* linear transcriptome. Because random primers were used to generate the sequencing library, the linear transcriptome not only contains mRNA, but all transcripts >150 nt, including non-coding transcripts. PCA revealed clear separation between high and low nutrient conditions along PC2 (13%), while Rhiz, Hn, and Ln/AM separated along PC1(23%), likely reflecting nitrogen availability (Figure 2A). DEG analysis for comparisons including rhizobia (Rhiz vs Hn, Rhiz vs Ln, Rhiz vs AM) had more upregulated DEGs, while non-Rhiz comparisons (Ln vs Hn and AM vs Hn) exhibited more downregulated DEGs (Figure 2B; Supplemental File 4). AM vs Ln treatments had only 11 DEGs, aligning with the limited sequencing depth and presumed early time point of colonization in AM samples. GO enrichment of DEGs revealed biological processes affected by symbiosis (Figure 2C). Comparisons involving rhizobia (Rhiz vs Hn, Rhiz vs Ln, and Rhiz vs AM) were enriched in cellular response to phosphate starvation, consistent with nitrogen-fixing symbiosis because nitrogen fixation is energetically demanding, requiring coordination with phosphate availability for ATP production and overall nutrient balance. In contrast, phosphate ion transport was the primary enriched process in AM vs Ln, suggesting a response to phosphate during early AM colonization. The AM vs Hn closely resembled Ln vs Hn, with enrichment in trehalose biosynthetic process, cell wall biogenesis, carbohydrate metabolic process, and ammonium transmembrane transport, highlighting the subtle but distinct transcriptional changes and possible early signaling of AM symbiosis.

**Figure 2.**
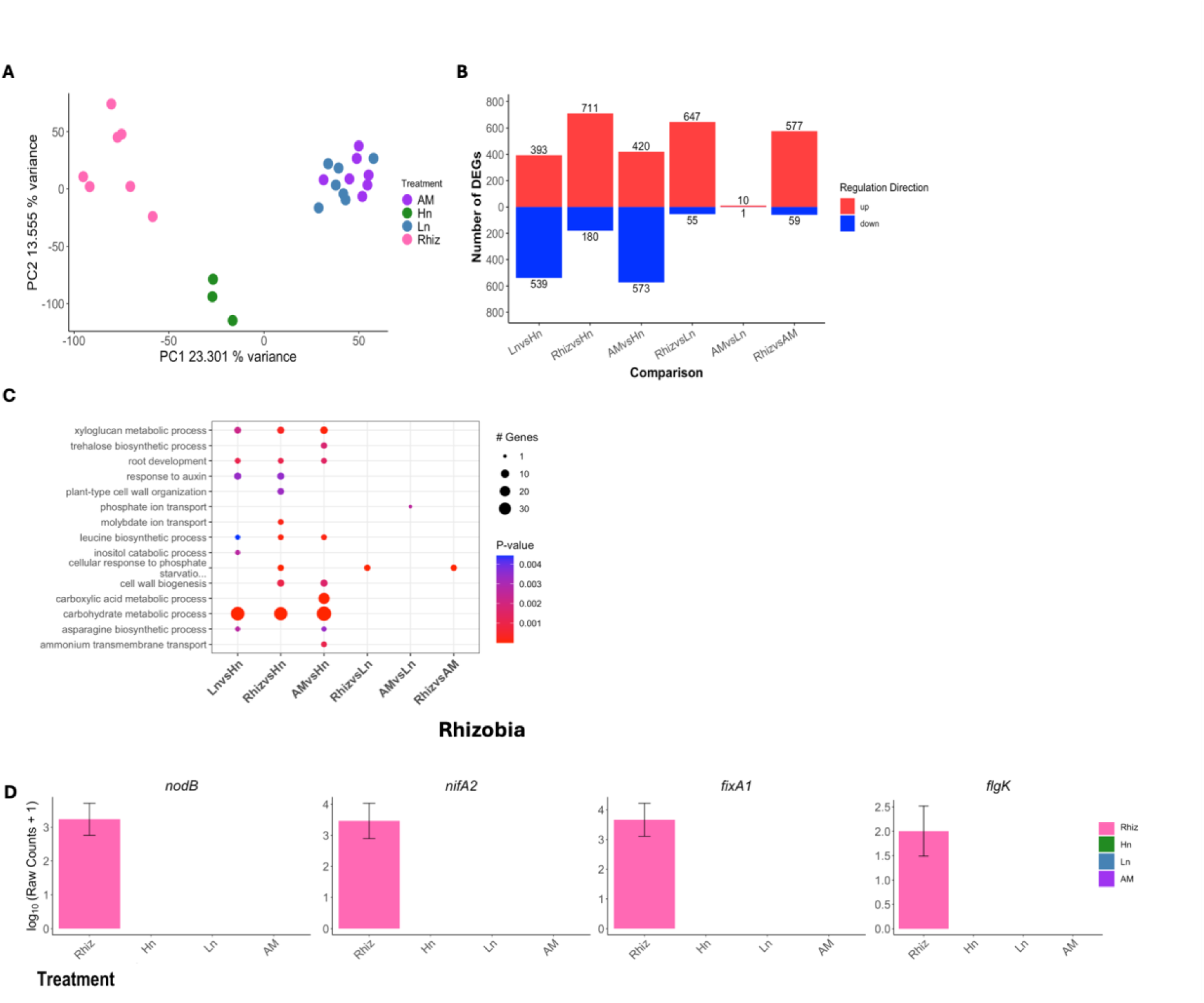
Transcriptomic analysis of *L. japonicus* and *M. loti*. (A) Principal component analysis (PCA) of linear-normalized counts across treatments. (B) Differentially expressed genes (DEGs) with FDR < 0.05 and |fold change| > 2, grouped by direction of regulation. (C) Representative Gene Ontology (GO) terms enriched among DEGs meeting the same criteria (See Supplemental File 4 for full list). (D–G) Expression profiles (log-transformed raw counts) of selected *M. loti* genes sampled from *L. japonicus* roots: (D) *nodB*, (E) *nifA2*, (F) *fixA1*, and (G) *flgK*. *nodB* and *flgK* are primarily expressed in the free-living state, while *nifA2* and *fixA1* are expressed in bacteroids. (D-G) Select *M. loti* gene expression is shown (log10(Raw Counts+1). Only treatments with mean raw counts ≥ 100 (mean raw counts ≥ 200 for *fixA1*) are displayed, highlighting that expression is largely restricted to Rhiz samples.

### 3.3 Microbial transcriptome

Deep sequencing of plant rRNA-depleted libraries constructed with random primers enabled simultaneous analysis of host and symbiont transcriptomes. While the sequencing depth and timepoint of harvest for the AM treatment were insufficient to detect transcripts beyond ribosomal genes, *M.loti* reads were sufficient for gene expression analysis. Most of the genes required for symbiosis in *M. loti* (strain R7A) are contained in a 502kb “symbiosis island”, which contains 414 genes. This region contains all the genes required for Nod factor synthesis (nod), nitrogen fixation (fix, nif), regulatory genes, transport genes, and an array of metabolic genes (Sullivan et al., 2001). In free-living bacteria, nodulation (nod) genes are expressed in response to plant-derived flavonoids or isoflavonoids for the biosynthesis of Lipochitooligosaccharides (LCOs), the bacteria-derived signaling molecules that bind to the LCO receptors on the plant membrane to signal symbiotic compatibility. The backbone of the chitooligosaccharide is synthesized by an N-acyltransferase (nodA), an N-deacetylase (nodB), and the chitin synthase (nodC). Other nod genes are involved in modifying the backbone through acylation (nodE and F), methylation (nodS), acetylation (nodL), or fucosylation (nodZ). The nodulation efficiency gene (nfeD) encodes a stomatin-like protein responsible for the bacterium’s competitiveness with the root (Green et al., 2009; A. Singh et al., 2021; Thomas & Rahman, 2020). We found expression of multiple nod genes in the *M. loti* inoculated samples (Figure 2D; Supplemental Figure 3A), indicating that the transcriptome data are from both free-living bacteria as well as symbiotic differentiated bacteroids.

Once symbiosis is established, genes required for N_2_-fixation are expressed. The *nif* operon is regulated by the NifA-RpoN-FixV cascade, with *nifA2* being essential for symbiotic nitrogen fixation in *L. japonicus*. We found expression of both *nifA1* and *nifA2* (Figure 2E; Supplemental Figure 3), confirming active nitrogen fixation pathways in our samples. The two *nifA* genes are not functionally redundant because a *nifA2* mutant forms ineffective nodules (Sullivan et al., 2001). In addition to *nif* genes, we observed expression of multiple *fix* genes, essential for nitrogen fixation in bacteroids (Figure 2F; Supplemental Figure 3). Genes encoding flagella and pili structures, such as the *flgK* gene, mostly found in free-living bacteria (Tatsukami et al., 2013), were also highly expressed (Figure 2G; Supplemental Figure 3). Samples likely contained a mixture of free-living bacteria and symbiotic bacterioids with *L. japonicus* nodules. Expression data for all *M. loti* genes are available in Supplemental File 5.

### 3.4 *L. japonicus* circRNA landscape

We identified circRNAs using two bioinformatic tools, CIRI2 and CircExplorer3 (CLEAR), to compensate for the high rate of false-positive discoveries and differences in the BSJ discovery algorithms (Budnick et al., 2024; Nielsen et al., 2022). One important difference is that CLEAR has no cutoff for the detection of BSJ reads, while CIRI2 requires >1 read to report a BSJ. To account for the differences in BSJ assignment, circRNAs with similar BSJs (Manhattan distance ≤ 10 across start and end coordinates) were clustered. A total of 15,252 unique nuclear circRNAs from 6,102 unique cognate gene IDs were predicted using the combined data set (Supplemental File 6). CLEAR and CIRI2 detected 12,779 (8,614 > 1 read) and 4,728 unique circRNAs, respectively. Of the 12,779 and 4,728 unique circRNAs found in CLEAR and CIRI2, 10,524 are only found in CLEAR and 2,473 are only found in CIRI2 (Supplemental Figure 4A). In CLEAR, there are 5,605 unique gene IDs, of which 3,660 are specific to CLEAR, and in CIRI2, there are 2,442 unique gene IDs, of which 497 are unique to CIRI2 (Supplemental Figure 4B). Populations of circRNAs from both CLEAR and CIRI2 had roughly 1/4th of circRNAs originating from intronic or intergenic regions (Supplemental Figure 4C). Due to our focus on coding RNAs, we have excluded intergenic circRNAs from all downstream analyses. However, we recognized the growing importance and relevance of lncRNA and analyzed whether there was a difference in the amount of unique intergenic circRNAs being detected between the treatments (Supplemental Figure 4D). The rhizobia treatment had the highest number of intergenic circRNAs. A very small number of circRNAs were identified as plastid (chloroplast and mitochondrial) from both detection tools, which will also be excluded from our analysis from henceforth, but are included in Supplemental File 6 (Supplemental Figure 4E). The average circRNA length for CLEAR was 1 kb, while the average length for CIRI2 is 800bp (Supplemental Figure 4F). CIRI2 was also observed to have a wider range of circRNA lengths versus CLEAR.

The majority of circRNAs identified have between 1 and 19 BSJ reads (Supplemental Figure 4G). CIRI2 identifies more circRNAs with higher read numbers. For instance, there are 149 circRNAs with greater than 81 reads, while CLEAR only has 45 circRNAs with a read number greater than 81 (Supplemental Figure 4G).

To investigate whether differences in sequencing depth might account for differences in detection or lack of detection of circRNA, we compared our read depths to the number of total circRNA reads for each sample from both analysis pipelines. Results show a strong correlation between the number of reads and the number of unique circRNAs identified for each pipeline (Supplemental Figure 5).

### 3.5 Specific circRNAs are found in association with symbionts

To explore the qualitative and quantitative distribution of nuclear genic circRNAs across treatments, we analyzed circRNAs identified by both CLEAR and CIRI2. As expected, fewer circRNAs were discovered in the AM treatment due to shallow sequencing (Supplemental Figure 6A). Among the other three treatments, Hn had the lowest number of circRNAs (n=3), while Ln and Rhiz showed higher circRNA abundance (Supplemental Figure 6A). When considering the average number of unique circRNAs from each treatment, the two treatments with increased nitrogen availability (Rhiz and Ln) had the highest numbers of both circRNAs and cognate genes (Supplemental Figure 6B and C). We also identified circRNAs that were specific to symbiont-treated samples: 4,290 circRNAs were unique to the Rhiz treatment, 1,491 were unique to the AM treatment, and 333 were shared between Rhiz and AM. In contrast, 1,059 circRNAs were shared among all treatments, and 62 circRNAs were present in every replicate of each treatment (Supplemental Figure 6D; Supplemental File 7).

Focusing on circRNAs conserved within treatments (present in ≥ 4 replicates, or ≥ 2 for Hn), we found 1,073 in Hn, 558 in Ln, 745 in Rhiz, and 295 in AM (Figure 3). Comparing these conserved circRNAs resulted in 221 shared among all treatments, while others were treatment-specific (Figure 3). Notably, there were 26 circRNAs specific to Rhiz, while none were exclusive to AM; two were found at the intersection of conserved AM and all Rhiz.

**Figure 3.**
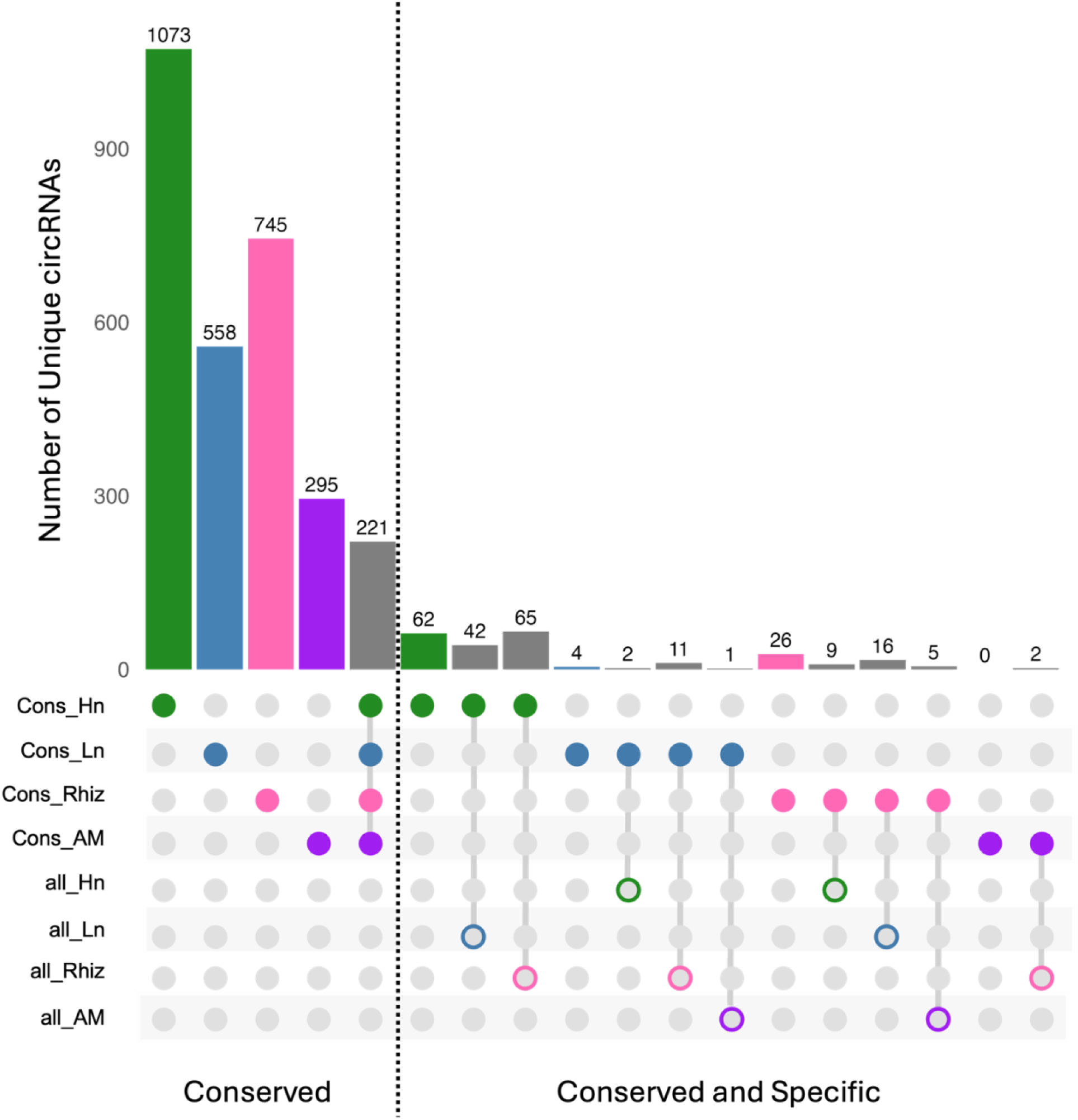
Upset plot of conserved and specific circRNAs. The left side displays circRNAs conserved within a single treatment (Cons_<treatment>), defined as present in ≥4 replicates for Ln, Rhiz, and AM, or ≥2 replicates for Hn. The right side shows overlaps of conserved circRNAs across treatments compared to all circRNAs from other treatments. Filled circles indicate that a circRNA meets the conservation threshold in that treatment, while open circles indicate that a circRNA is present in at least one replicate of that treatment. Intersections of circles across treatments show circRNAs shared between those treatments. For example, comparing the 1,073 circRNAs from cons_Hn against all circRNAs in Ln (all_Ln) and Rhiz (all_Rhiz) leaves 62 specific to cons_Hn, 42 shared between cons_Hn and Ln, and 65 shared between cons_Hn and Rhiz. Full lists of circRNAs are provided in Supplemental File 7.

To highlight potentially relevant circRNAs, we identified several with treatment-specific or shared patterns across biologically meaningful conditions (Supplemental Table 1). For example, a circRNA from LotjaGi1g1v0118200 (chr1:18680735–18681182), encoding Asparagine Synthetase, was detected at the intersection of conserved Hn and all Ln (represented by the number 42 in Figure 3)—treatments that differ in nitrate levels in the growth media. Another circRNA from the same gene family (chr6:36811083–36811521) was found in the intersection of conserved Rhiz and all Hn (represented by the number 9 in Figure 3), again supporting a nitrogen-related role.

In the Rhiz treatment, a circRNA from LotjaGi5g1v0169900, annotated as Type I inositol-1,4,5-trisphosphate 5-phosphatase, stood out due to its high expression and potential involvement in phosphate signaling, a pathway known to influence rhizobial symbiosis (Riemer et al., 2022).

Finally, we detected a circRNA originating from *SymRK* (LotjaGi2g1v0330500, chr2:80616495–80617260), a key receptor kinase in symbiosis signaling, in AM and Rhiz treatments. Although not present in all replicates, its specific occurrence in both symbiotic treatments points to a potential regulatory role. The absence in some replicates highlights the inherent variability and detection sensitivity of circRNA discovery methods. Additional treatment-specific circRNAs of interest are listed in Supplemental Table 1, with the full set of conserved and intersecting circRNAs available in Supplemental File 7.

### 3.6 Symbiotic Signaling and the Common Symbiosis Pathway

To explore transcriptional regulation within symbiotic signaling, we examined differentially expressed genes (DEGs) across treatment comparisons. Notably, *CLE-RS2* and *CLE-RS3* exhibited dramatic upregulation in Rhiz vs. Ln and Rhiz vs. AM comparisons (logFC ∼10), while being significantly downregulated in Ln vs. Hn, Rhiz vs. Hn, and AM vs. Hn contrasts. A similar expression pattern was observed for *NIN* and *TML*, genes known to function in nodule organogenesis and autoregulation of nodulation, respectively (Soyano et al., 2014; Tsikou et al., 2018) (Supplemental Figure 7).

To explore post-transcriptional regulation in the CSP pathway, we identified circRNAs associated with CSP genes. We detected circRNAs from 38 of the 61 known/predicted symbiotic-related genes, highlighting their potential regulatory role in symbiosis (Figure 4; Supplemental Figure 8; Supplemental File 6).

**Figure 4.**
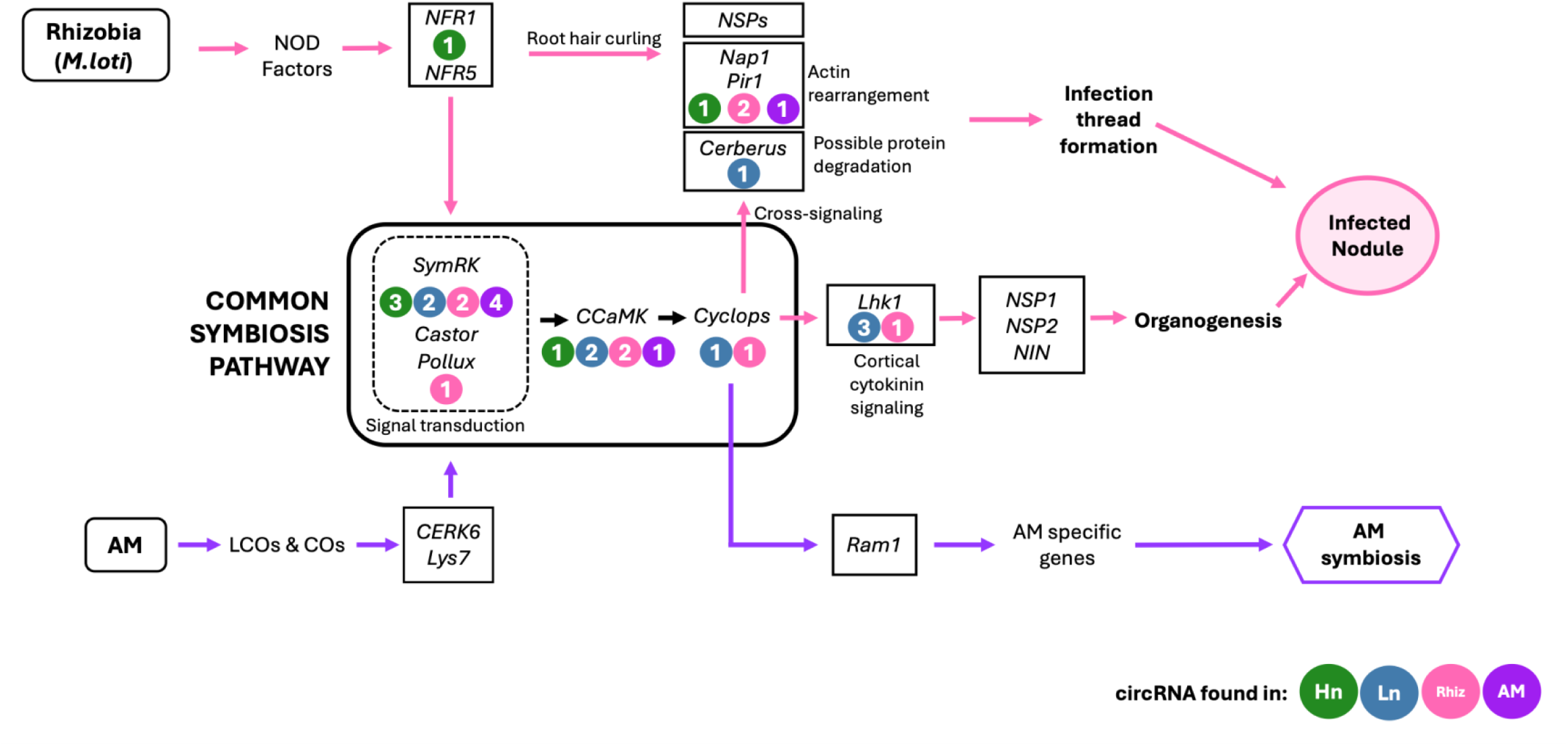
The Common Symbiotic Pathway. Schematic of the CSP pathway showing which genes were identified to have a circRNA. The color of the circle corresponds to the treatment, and the number inside the circle represents how many circRNAs from that treatment were found.

In the early perception stage, one circRNA was found in the Hn treatment associated with Nod factor/LCO receptors. In the infection thread pathway, three circRNAs were identified for *Pir1* (actin rearrangement): one specific to Rhiz, one to AM, and one shared between Rhiz and Hn. Three circRNAs were also detected for *Cerberus*, all in Ln. In the CSP pathway, *SymRK* had nine circRNAs: two specific to Hn, two to Ln, and three to AM, with two others shared across treatments. *Pollux* had one Rhiz-specific circRNA. *CCaMK* had four circRNAs, with two being treatment-specific (Rhiz and Hn) and two shared across treatments (AM/Rhiz and Ln/Hn). *Cyclops* had one circRNA shared between Ln and Rhiz. *SymRK, Pollux, CCaMK,* and *Cyclops* are also recognized in the establishment of AM. Continuing with the organogenesis pathway in the establishment of nodules, *Lhk1* had five circRNAs: two each in Rhiz and Ln, and one shared. Notably, its paralog *Lhk1a* had 16 circRNAs that were detected in all treatments, though in varying numbers. Identification of these differences in circRNA expression from symbiosis genes suggests that there may be an additional transcriptional regulatory layer within the symbiotic signaling network during symbiotic interactions.

### 3.7 Validation

Validation is essential after the identification of circRNAs via different bioinformatic tools because of the use of different aligners and the chance of false positives. RT-PCR using divergent primers was used to validate putative potential circRNAs. Unlike the typical convergent primers used to amplify linear cDNA fragments, divergent primers are oriented away from each other and cannot amplify linear RNA but capture the BSJ in circular RNA and/or the internal sequence (Figure 5A). As a control, gDNA is used to ensure there is no tandem duplication in the genome.

**Figure 5.**
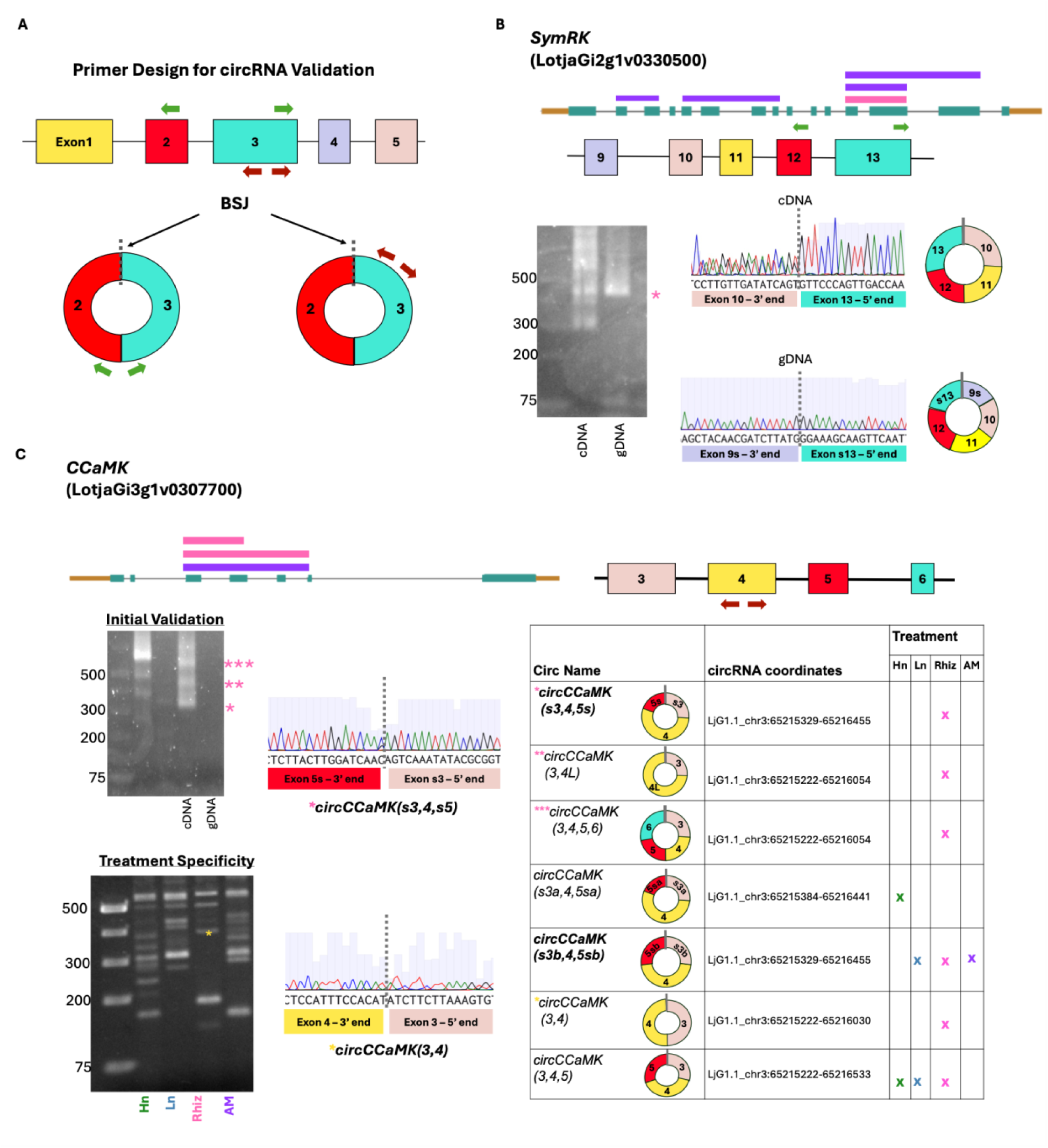
Validation of *SymRK* and *CCaMK* circRNAs. (A) Schematic of divergent primer design for circRNA validation. Primers are designed to capture the BSJ (green arrows) and/or to capture the internal sequence, thereby maximizing the potential for capturing multiple circRNAs with a shared exon (red arrows). (B) Evidence of false-positive *SymRK* circRNA. (C) Validation of *CCaMK* circRNAs. Initial validation for *SymRK* and *CCaMK* was completed using Rhiz tissue. Purple and pink bars indicate circRNAs detected in sequencing. Full image of gels (Supplemental Figures 11 and 12). All coordinates of validated circRNAs can be found in Supplemental File 2.

We successfully validated five circRNAs from several genes using *Rhiz* treatment tissue (Supplemental Figure 9). Notably, circCCR4(4,5), previously confirmed in leaf tissue (Budnick et al., 2024) and derived from *Cinnamoyl-CoA reductase 4* (*CCR4*; *LotjaGi1g1v0115700*), was also validated in root tissue. Here, however, we focus on circRNAs detected in Rhiz and AM treatments from two core CSP genes: *SymRK* and *CCaMK*.

For *SymRK,* five circRNAs were identified from Rhiz and AM treatments: one circRNA from Rhiz, three circRNAs from AM, and one circRNA that was shared between both AM and Rhiz treatments (Figure 5B). We attempted to validate the shared circRNA (LjG1.1_chr2:80616495-80617260) using Rhiz tissue. However, an identical band appeared in the gDNA, indicating a false-positive circRNA resulting from a tandem duplication (Figure 5B). Despite a slight mismatch in the BSJ between the cDNA and gDNA, the extensive sequence overlap led us to classify it as a false-positive.

Two different circRNAs were identified for *CCaMK*: one specific to Rhiz and the other shared between AM and Rhiz (Figure 5C). Initial validation using Rhiz tissue with primers targeting exon 4 confirmed three novel circRNAs that were not detected by sequencing. Expanding validation to other treatments revealed tissue specificity. First, *circCCaMK(3,4,5)* was present in Hn, Ln, and Rhiz, but absent in AM. Second, two nearly identical circRNAs, *circCCaMK(s3a,4,5sa)* in Hn and *circCCaMK(s3b,4,5sb)* in Rhiz, Ln, and AM, shared exon composition but had slightly different BSJs. Lastly, we validated a Rhiz-specific circRNA, *circCCaMK(3,4)* (Figure 5C; Supplemental Figure 10). This validation demonstrates that expression of circRNAs from the CcaMK locus is at least qualitatively different across symbiosis treatment conditions.

### 3.8 MicroRNA Recognition Elements and competing endogenous RNA networks

MiRNAs can bind to circRNAs, competing with their cognate linear mRNAs and potentially reducing mRNA degradation or enhancing translation (Figure 6A) (Salmena et al., 2011). To identify putative miRNA binding sites, or miRNA recognition elements (MREs), we reconstructed full-length circRNA sequences and analyzed them with psRNATarget v2 (Dai & Zhao, 2011) using *L. japonicus* miRNAs from miRBase (Kozomara & Griffiths-Jones, 2014) (see Figure 2 and Methods for details).

**Figure 6.**
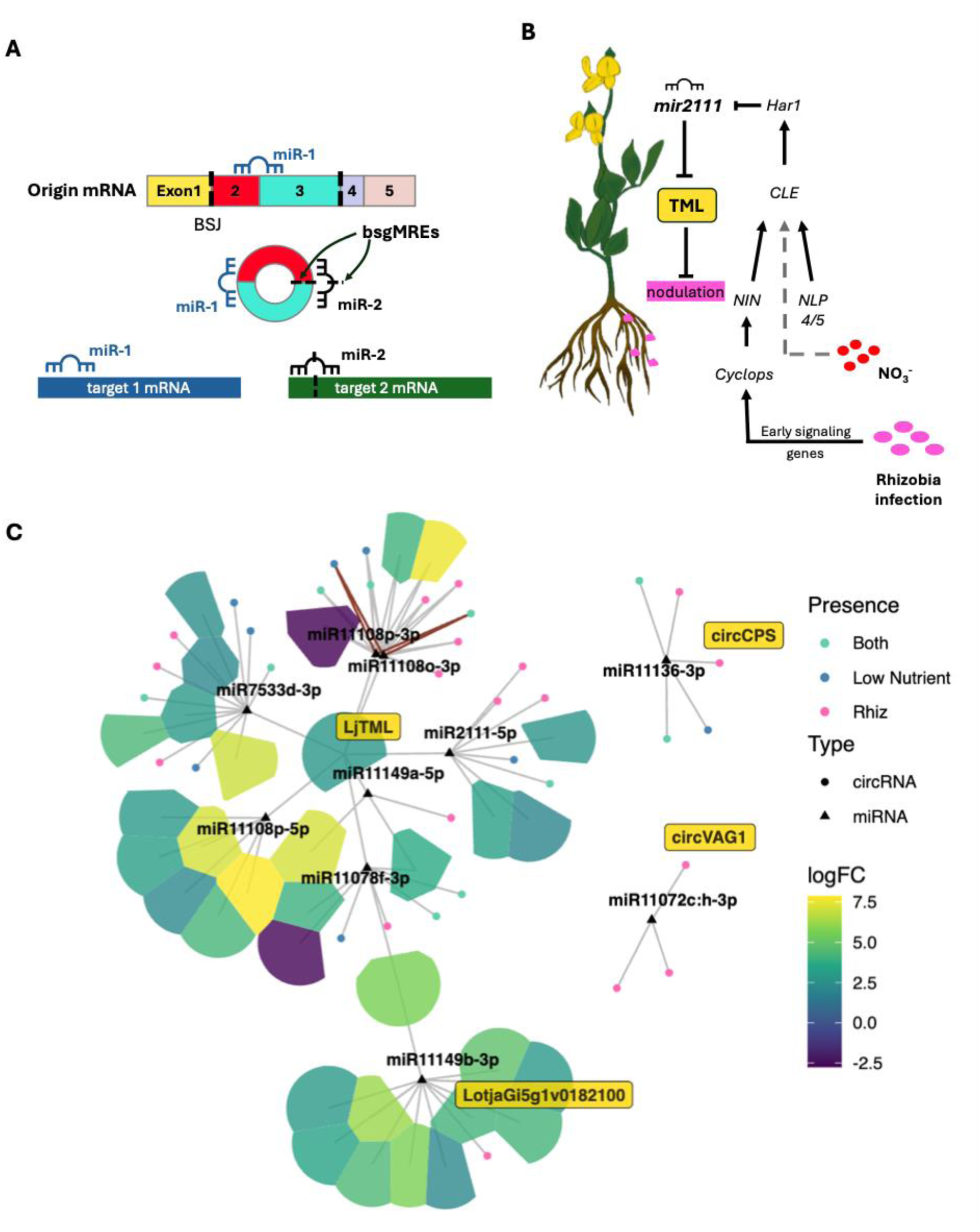
Backsplice-generated MREs, mir2111, and TML focused ceRNA network. (A) Novel MREs can be generated by the BSJ (bsgMREs) and thereby interact with different miRNAs that are not targeting the cognate RNA, but a different network of target RNAs. (B) Schematic representation of the autoregulation of nodulation (AON) pathway. (C) ceRNA network with a focus on TML in the Rhiz vs Ln comparison. Nodes labeled with gold background are CSP genes or circRNAs from a CSP parent gene. Voronoi tile nodes represent DEGs colored by their logFC, with a positive logFC indicating higher expression in Rhizobia compared to Low Nutrient treatments. Circular circRNA isoform nodes are colored by presence in either Rhizobia, Low Nutrient, or Both treatments. Triangular miRNA nodes are labeled, some miRNAs with identical targets in the subnetwork were collapsed, and the node was given a name to represent all of them (for example, miR11072c:h is a collapsed node including 6 isoforms). Brown lines represent circRNAs containing bsgMREs.

We identified 2,374 unique circRNAs with MREs to 362 of the 365 annotated *L. japonicus* miRNAs (Supplemental Figure 4; Supplemental File 8). Most circRNAs contain one MRE, while the majority of circRNAs do not contain a MRE (Supplementary Figure 13). Comparing MRE-containing circRNAs across treatments, we found eight that were present in at least three Ln (136 circRNAs), three Rhiz (165), two Hn (185), and any AM (12) samples, and considered them conserved (Supplemental Figure 14A; Supplemental Table 2).

To identify treatment-specific circRNAs with MREs, we compared those present in at least three Ln or Rhiz samples (out of 7) and at least two Hn samples (out of 3) against circRNAs from other treatments. This yielded 1,191 MRE circRNAs in Rhiz, 1,195 in Ln, 864 in Hn, and only 12 in AM. The low number in AM and Hn likely reflects reduced sample numbers and sequencing depth, which limits CIRI-vis’s ability to reconstruct full-length circRNAs. We identified 15 Hn, 2 Ln, and 10 Rhiz-specific MRE circRNAs (Supplemental Figure 14B-D; Supplemental Table 3).

Theoretically, the BSJ could generate novel MREs, backsplice-generated MREs (bsgMREs), that are not present in cognate linear mRNAs (Figure 6A). We found 294 circRNAs containing bsgMREs targeting 216 unique miRNAs. Three of the Rhiz-specific circRNAs and one Ln-specific circRNA contained bsgMREs, while none of the Hn-specific circRNAs did (Supplemental Table 3). No bsgMRE circRNAs were conserved across all treatments, largely due to limited detection in AM, where only one bsgMRE circRNA was identified. Treatment-specific MREs are available in Supplemental Table 4.

An established example of a miRNA-mediated symbiosis regulation is mir2111, which represses its target gene *TOO MUCH LOVE (TML)*, a negative regulator of nodulation (Figure 6B) (Tsikou et al., 2018). Under low nitrogen conditions, mir2111 is upregulated and moves through the phloem to suppress TML, enabling nodulation. Conversely, when nitrogen is sufficient or rhizobia colonization is extensive, mir2111 levels drop, allowing TML to limit further nodulation (Tsikou et al., 2018). While we did not sequence miRNAs directly, we observed expression of several key components of the autoregulation of nodulation (AON) pathway in which miR2111 operates, including *NIN*, *TML*, and CLE peptide genes (CLE-RS2 and CLE-RS3), which collectively help balance nodulation in response to rhizobia signals and nitrate availability (see Section 3.6; Supplemental Figure 7). The role of mir2111 highlights how miRNAs can act as key regulators in symbiosis by repressing their targets. Beyond individual reactions, the competing endogenous RNA (ceRNA) hypothesis suggests that different RNA molecules, including mRNAs, long non-coding RNAs, and circRNAs, can regulate each other by competing for shared miRNAs (Salmena et al., 2011).

To investigate this hypothesis, we constructed a network linking circRNA:miRNA:DEG predictions to better understand ceRNAs’ interactions in the autoregulation of nodulation during symbiotic signaling (Figure 6C). For the creation of the symbiotic signaling subnetwork, three nodes were highlighted based on genes involved in autoregulation of nodulation and other symbiotic signaling genes: repressor of nodulation, LjTML, a circRNA from LjVAG1 (LotjaGi2g1v0179500_LC), a gene important to endoreduplication in growing nodules (Suzaki et al., 2014), and a circRNA from LjCPS (LotjaGi1g1v0228700), a gene known for its role in synthesizing plant hormones like gibberellins (Maekawa et al., 2009). Nodes within two connections of these were included. The predicted circRNA:miRNA:DEG network shows how much competition for the miRNA binding to circRNA versus mRNA is present at the post-transcriptional level, which may play a role in symbiosis.

For example, consider the connection between miR2111 and TML. In the Rhiz vs. Ln comparison, miR2111 also targets three other DEGs. Five circRNAs contain a predicted MRE for miR2111, three of which are isoform-specific to the Rhiz treatment. These circRNAs originate from three different genes. Additional details about these ceRNAs can be found in Table 1.

**Table 1.**
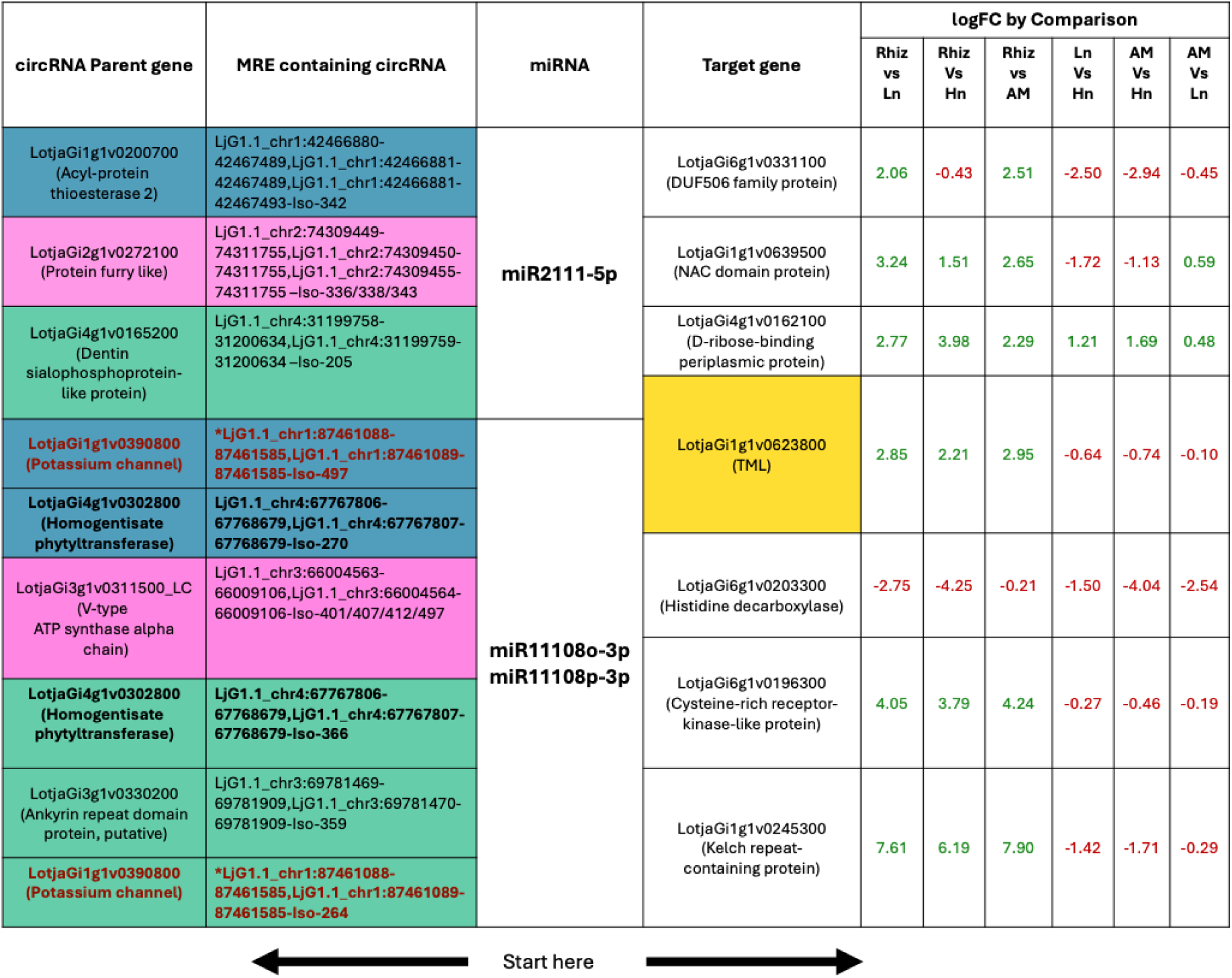
Examples of two miRNAs and their potential targets from the ceRNA network. miR2111 is the node to the right above TML in Figure 7. The three circs from Rhizobia are isoforms of the same circRNA. mir11108o/p-3p is the node directly above TML in Figure 7. Isoforms have been collapsed. Colored font for circRNA represents those circRNAs with a bsgMRE; circRNAs in bold originate from the same parent gene. TML is in gold.

The presence of circCPS and circVAG1 (highlighted in yellow in Figure 6C) and their interactions with their respective miRNAs suggest that some symbiosis-related circRNAs may function as miRNA sponges to regulate gene expression. For example, CircVAG1 harbors an MRE for six isoforms of miR11072c:h-3p.

Notably, this network reveals the presence of bsgMRE circRNAs, indicated by brown lines in Figure 7C and brown and bold text in Table 1. Two isoforms of the same miRNA (miRNAs originating from different genes) can bind to two isoforms of the same circRNA (circRNAs with the same BSJ, different internal sequence), one isoform found in Ln only, and the other isoform was found in both Rhiz and Ln. This binding is predicted based on the hypothesis that the BSJ can generate a novel MRE (bsgMRE), distinct from the linear sequence. These elements highlight the complexity of post-transcriptional regulation in symbiosis.

## 4 Discussion

In this study, we investigated the circRNA landscape and transcriptome of *Lotus japonicus* (Gifu) under symbiotic conditions with arbuscular mycorrhizal (AM) fungi or rhizobia (Rhiz), alongside nutrient treatments with high or low N and P in the media. Analysis of the linear transcriptomes revealed gene expression patterns of our host as well as rhizobia, as bacteroids or free-living bacteria. We focused on the identification of circRNAs, particularly those involved in well-known symbiotic pathways, and we explored interactions of identified circRNAs, linear DEGs, and predicted miRNAs in ceRNA networks. Additionally, we show that BSJ can generate novel miRNA recognition elements (bsgMREs) that could compete with a unique set of miRNA interactions beyond those of their linear transcripts. Lastly, we validated several circRNAs originating from symbiosis and metabolism-related genes.

Circular RNAs are only distinguishable from their linear counterparts by their backsplice junctions (BSJs), making accurate detection technically challenging. Genetic features like repetitive sequences, genomic rearrangements, experimental variability, and technical artifacts during extraction, reverse transcription, sequencing, and bioinformatic analysis can affect the accurate detection of the BSJ, leading to false positives (Budnick et al., 2024; Hansen et al., 2016; Utley et al., 2025). Additionally, low sequencing depth, particularly in our AM samples, can limit detection sensitivity and bias downstream analyses.

To mitigate these issues, various bioinformatic pipelines have been developed to improve circRNA detection and sensitivity. A recent benchmarking study showed that while circRNA detection tools generally demonstrate high precision (over 95%), they vary widely in sensitivity, by as much as 50-fold (Vromman et al., 2023). No single approach identified all true circRNAs (X. Zeng et al., 2017). In our study, we employed both CIRI2 (Gao et al., 2018) and CLEAR (X.-K. Ma et al., 2019), which are known for their high precision and relatively high sensitivity. This multi-bioinformatic pipeline strategy aligns with best practices in the field and reduces the likelihood of false positives (Budnick et al., 2024; Hansen et al., 2016; Utley et al., 2025).

Beyond computational work, experimental validation is essential for determining true circRNAs and their function. In this study, we validated 12 circRNAs derived from conserved and treatment-specific genes, including those involved in lipid metabolism and the Common Symbiosis Pathway (CSP) (Figure 5, 6, Supplemental Figure 9). For example, the Rhiz-specific circCCaMK(3,4) could play a role in symbiotic regulation, while a putative circRNA from SymRK was found to be a genomic duplication (Figure 5). These results emphasize the importance of experimental validation to validate true circRNAs.

Few plant studies have investigated circRNAs and their role in symbiosis. In tomato, 146 circRNAs were found to respond to AM fungi inoculation, including 88 up-regulated and 58 down-regulated circRNAs (Z. Zeng et al., 2023). In *Phaseolus vulgaris*, 3,448 circRNAs were expressed during nodulation, with the highest numbers in the nitrogen-fixing stage (Wu et al., 2020). In *Medicago truncatula*, 19 circRNAs were identified under severe drought stress in the presence of rhizobia, pointing to circRNAs’ potential role in coordinating and regulating abiotic and biotic signaling (Jing et al., 2022).

Our study builds on this work by focusing on circRNA originating from CSP genes, which are essential for both AM and rhizobia symbiosis. For example, the work in *Phaseolus vulgaris* identified circRNAs from CSP genes during specific stages of symbiosis (Wu et al., 2020). While previous studies explore ceRNA networks in the context of symbiosis, we expand on this concept by incorporating backsplice-generated miRNA recognition elements (bsgMREs), which are novel recognition sites created by the backsplice junction that are absent from linear transcripts.

We placed particular focus on the *TML*-*mir2111* interaction, a well-characterized component of the Autoregulation of Nodulation (AON) pathway in *L. japonicius*, which regulates nodulation through long-distance signaling (Okuma & Kawaguchi, 2021; Tsikou et al., 2018). While we did not identify circRNAs from AON pathway genes, we were able to predict interactions between predicted miRNAs, circRNAs, and DEGs in Rhiz vs Ln treatments (Figure 6C). Notably, including two bsgMRE circRNAs that interact with different isoforms of the same miRNA, highlighting an example of the levels of regulatory complexity. These findings support the idea that circRNAs contribute to ceRNA networks in symbiosis.

Moreover, the mobility of mir2111, known to travel from shoot to root to regulate nodulation via TML, raises the possibility that circRNAs may also be mobile. Especially due to the presence of circRNAs in exosomes (Hansen et al., 2013), circRNAs may be able to function beyond their tissue of origin, making implications for function in systemic or even plant-microbe communication. Additional evidence suggests that circRNAs participate in transcriptional regulation via R-loop (RNA:DNA hybrid) formation (Conn et al., 2017) or are translated into peptides (Pan et al., 2025).

Future studies will focus on validating circRNAs from key CSP genes to confirm the regulatory role in symbiosis and nutrient stress. High-throughput approaches, such as the circPanel method (Budnick et al., 2024; Rahimi et al., 2021), could streamline this process. Integration of deep circular, linear, and miRNA sequencing, normalized with spike-ins (Laosuntisuk et al., 2024), would provide a more accurate representation of ceRNA networks. Sampling across nodulation stages and using symbiotic mutants such as *ccamk* (Gleason et al., 2006) and *snf* (Tirichine et al., 2006) and others could further clarify the temporal and genetic regulation of circRNAs. Such work would build on our findings and deepen understanding of post-transcriptional regulation in *L. japonicus* symbiosis.

## References

Adrian Alexa, J. R. (2017). *topGO* [Computer software]. Bioconductor. 10.18129/B9.BIOC.TOPGO

Andrews, S. (2010). Babraham Bioinformatics—FastQC A Quality Control tool for High Throughput Sequence Data. https://www.bioinformatics.babraham.ac.uk/projects/fastqc/

Bolger, A. M., Lohse, M., & Usadel, B. (2014). Trimmomatic: A flexible trimmer for Illumina sequence data. Bioinformatics, 30(15), 2114–2120. 10.1093/bioinformatics/btu170

Broghammer, A., Krusell, L., Blaise, M., Sauer, J., Sullivan, J. T., Maolanon, N., Vinther, M., Lorentzen, A., Madsen, E. B., Jensen, K. J., Roepstorff, P., Thirup, S., Ronson, C. W., Thygesen, M. B., & Stougaard, J. (2012). Legume receptors perceive the rhizobial lipochitin oligosaccharide signal molecules by direct binding. Proceedings of the National Academy of Sciences, 109(34), 13859–13864. 10.1073/pnas.1205171109

Budnick, A., Franklin, M. J., Utley, D., Edwards, B., Charles, M., Hornstein, E. D., & Sederoff, H. (2024). Long- and short-read sequencing methods discover distinct circular RNA pools in Lotus japonicus. The Plant Genome, e20429. 10.1002/tpg2.20429

Bushnell, B. (2014). BBMap: A Fast, Accurate, Splice-Aware Aligner. https://escholarship.org/uc/item/1h3515gn

Chang, S., Puryear, J., & Cairney, J. (1993). A simple and efficient method for isolating RNA from pine trees. Plant Molecular Biology Reporter, 11(2), Article 2. 10.1007/BF02670468

Charpentier, M., Bredemeier, R., Wanner, G., Takeda, N., Schleiff, E., & Parniske, M. (2008). Lotus japonicus CASTOR and POLLUX Are Ion Channels Essential for Perinuclear Calcium Spiking in Legume Root Endosymbiosis. The Plant Cell, 20(12), 3467–3479. 10.1105/tpc.108.063255

Conn, V. M., Hugouvieux, V., Nayak, A., Conos, S. A., Capovilla, G., Cildir, G., Jourdain, A., Tergaonkar, V., Schmid, M., Zubieta, C., & Conn, S. J. (2017). A circRNA from SEPALLATA3 regulates splicing of its cognate mRNA through R-loop formation. Nature Plants, 3(5), 1–5. 10.1038/nplants.2017.53

Dai, X., & Zhao, P. X. (2011). psRNATarget: A plant small RNA target analysis server. Nucleic Acids Research, 39(suppl_2), W155–W159. 10.1093/nar/gkr319

De Luis, A., Markmann, K., Cognat, V., Holt, D. B., Charpentier, M., Parniske, M., Stougaard, J., & Voinnet, O. (2012). Two MicroRNAs linked to nodule infection and nitrogen-fixing ability in the legume Lotus japonicus. Plant Physiology, 160(4), 2137–2154. 10.1104/pp.112.204883

Fan, J., Quan, W., Li, G.-B., Hu, X.-H., Wang, Q., Wang, H., Li, X.-P., Luo, X., Feng, Q., Hu, Z.-J., Feng, H., Pu, M., Zhao, J.-Q., Huang, Y.-Y., Li, Y., Zhang, Y., & Wang, W.-M. (2020). circRNAs Are Involved in the Rice-Magnaporthe oryzae Interaction1 [OPEN]. Plant Physiology, 182(1), 272–286. 10.1104/pp.19.00716

Fernández, I., Cosme, M., Stringlis, I. A., Yu, K., de Jonge, R., van Wees, S. C. M., Pozo, M. J., Pieterse, C. M. J., & van der Heijden, M. G. A. (2019). Molecular dialogue between arbuscular mycorrhizal fungi and the nonhost plant Arabidopsis thaliana switches from initial detection to antagonism. New Phytologist, 223(2), 867–881. 10.1111/nph.15798

Gao, Y., Zhang, J., & Zhao, F. (2018). Circular RNA identification based on multiple seed matching. Briefings in Bioinformatics, 19(5), 803–810. 10.1093/bib/bbx014

Gleason, C., Chaudhuri, S., Yang, T., Muñoz, A., Poovaiah, B. W., & Oldroyd, G. E. D. (2006). Nodulation independent of rhizobia induced by a calcium-activated kinase lacking autoinhibition. Nature, 441(7097), 1149–1152. 10.1038/nature04812

Green, J. B., Lower, R. P. J., & Young, J. P. W. (2009). The NfeD Protein Family and Its Conserved Gene Neighbours Throughout Prokaryotes: Functional Implications for Stomatin-Like Proteins. Journal of Molecular Evolution, 69(6), 657–667. 10.1007/s00239-009-9304-8

Guria, A., Sharma, P., Natesan, S., & Pandi, G. (2020). Circular RNAs—The Road Less Traveled. Frontiers in Molecular Biosciences, 6. 10.3389/fmolb.2019.00146

Hansen, T. B., Jensen, T. I., Clausen, B. H., Bramsen, J. B., Finsen, B., Damgaard, C. K., & Kjems, J. (2013). Natural RNA circles function as efficient microRNA sponges. Nature, 495(7441), 384–388. 10.1038/nature11993

Hansen, T. B., Venø, M. T., Damgaard, C. K., & Kjems, J. (2016). Comparison of circular RNA prediction tools. Nucleic Acids Research, 44(6), e58. 10.1093/nar/gkv1458

Hornstein, E. D., Charles, M., Franklin, M., Edwards, B., Vintila, S., Kleiner, M., & Sederoff, H. (2024). IPD3, a master regulator of arbuscular mycorrhizal symbiosis, affects genes for immunity and metabolism of non-host Arabidopsis when restored long after its evolutionary loss. Plant Molecular Biology, 114(2), 21. 10.1007/s11103-024-01422-3

Jing, J., Yang, P., Wang, Y., Qu, Q., An, J., Fu, B., Hu, X., Zhou, Y., Hu, T., & Cao, Y. (2022). Identification of Competing Endogenous RNAs (ceRNAs) Network Associated with Drought Tolerance in Medicago truncatula with Rhizobium Symbiosis. International Journal of Molecular Sciences, 23(22), Article 22. 10.3390/ijms232214237

Kefas, B., Godlewski, J., Comeau, L., Li, Y., Abounader, R., Hawkinson, M., Lee, J., Fine, H., Chiocca, E. A., Lawler, S., & Purow, B. (2008). microRNA-7 Inhibits the Epidermal Growth Factor Receptor and the Akt Pathway and Is Down-regulated in Glioblastoma. Cancer Research, 68(10), 3566–3572. 10.1158/0008-5472.CAN-07-6639

Kozomara, A., & Griffiths-Jones, S. (2014). miRBase: Annotating high confidence microRNAs using deep sequencing data. Nucleic Acids Research, 42(D1), D68–D73. 10.1093/nar/gkt1181

Laosuntisuk, K., Vennapusa, A., Somayanda, I. M., Leman, A. R., Jagadish, S. K., & Doherty, C. J. (2024). A normalization method that controls for total RNA abundance affects the identification of differentially expressed genes, revealing bias toward morning-expressed responses. The Plant Journal, 118(5), 1241–1257. 10.1111/tpj.16654

Li, Y., Zheng, F., Xiao, X., Xie, F., Tao, D., Huang, C., Liu, D., Wang, M., Wang, L., Zeng, F., & Jiang, G. (2017). CircHIPK3 sponges miR-558 to suppress heparanase expression in bladder cancer cells. EMBO Reports, 18(9), 1646–1659. 10.15252/embr.201643581

Liao, Y., Smyth, G. K., & Shi, W. (2014). featureCounts: An efficient general purpose program for assigning sequence reads to genomic features. Bioinformatics, 30(7), 923–930. 10.1093/bioinformatics/btt656

Liu, H., Yu, W., Wu, J., Li, Z., Li, H., Zhou, J., Hu, J., & Lu, Y. (2020). Identification and characterization of circular RNAs during wood formation of poplars in acclimation to low nitrogen availability. Planta, 251(2), 47. 10.1007/s00425-020-03338-w

Lv, L., Yu, K., Lü, H., Zhang, X., Liu, X., Sun, C., Xu, H., Zhang, J., He, X., & Zhang, D. (2020). Transcriptome-wide identification of novel circular RNAs in soybean in response to low-phosphorus stress. PLOS ONE, 15(1), e0227243. 10.1371/journal.pone.0227243

Ma, P., Gao, S., Zhang, H. y., Li, B. y., Zhong, H. x., Wang, Y. k., Hu, H. m., Zhang, H. k., Luo, B. w., Zhang, X., Liu, D., Wu, L., Gao, D. j., Gao, S. q., Zhang, S. z., & Gao, S. b. (2021). Identification and characterization of circRNAs in maize seedlings under deficient nitrogen. Plant Biology, 23(5), 850–860. 10.1111/plb.13280

Ma, X.-K., Wang, M.-R., Liu, C.-X., Dong, R., Carmichael, G. G., Chen, L.-L., & Yang, L. (2019). CIRCexplorer3: A CLEAR Pipeline for Direct Comparison of Circular and Linear RNA Expression. Genomics, Proteomics & Bioinformatics, 17(5), 511–521. 10.1016/j.gpb.2019.11.004

Maekawa, T., Maekawa-Yoshikawa, M., Takeda, N., Imaizumi-Anraku, H., Murooka, Y., & Hayashi, M. (2009). Gibberellin controls the nodulation signaling pathway in Lotus japonicus. The Plant Journal, 58(2), 183–194. 10.1111/j.1365-313X.2008.03774.x

Murakami, E., Cheng, J., Gysel, K., Bozsoki, Z., Kawaharada, Y., Hjuler, C. T., Sørensen, K. K., Tao, K., Kelly, S., Venice, F., Genre, A., Thygesen, M. B., Jong, N. de, Vinther, M., Jensen, D. B., Jensen, K. J., Blaise, M., Madsen, L. H., Andersen, K. R.,… Radutoiu, S. (2018). Epidermal LysM receptor ensures robust symbiotic signalling in Lotus japonicus. eLife, 7, e33506. 10.7554/eLife.33506

Nielsen, A. F., Bindereif, A., Bozzoni, I., Hanan, M., Hansen, T. B., Irimia, M., Kadener, S., Kristensen, L. S., Legnini, I., Morlando, M., Jarlstad Olesen, M. T., Pasterkamp, R. J., Preibisch, S., Rajewsky, N., Suenkel, C., & Kjems, J. (2022). Best practice standards for circular RNA research. Nature Methods, 19(10), 1208–1220. 10.1038/s41592-022-01487-2

Okuma, N., & Kawaguchi, M. (2021). Systemic Optimization of Legume Nodulation: A Shoot-Derived Regulator, miR2111. Frontiers in Plant Science, 12. 10.3389/fpls.2021.682486

Pan, X., Xu, S., Cao, G., Chen, S., Zhang, T., Yang, B. B., Zhou, G., & Yang, X. (2025). A novel peptide encoded by a rice circular RNA confers broad-spectrum disease resistance in rice plants. The New Phytologist. 10.1111/nph.70018

Pertea, M., Pertea, G. M., Antonescu, C. M., Chang, T.-C., Mendell, J. T., & Salzberg, S. L. (2015). StringTie enables improved reconstruction of a transcriptome from RNA-seq reads. Nature Biotechnology, 33(3), 290–295. 10.1038/nbt.3122

Rahimi, K., Venø, M. T., Dupont, D. M., & Kjems, J. (2021). Nanopore sequencing of brain-derived full-length circRNAs reveals circRNA-specific exon usage, intron retention and microexons. Nature Communications, 12(1), 4825. 10.1038/s41467-021-24975-z

Ren, B., Wang, X., Duan, J., & Ma, J. (2019). Rhizobial tRNA-derived small RNAs are signal molecules regulating plant nodulation. Science, 365(6456), 919–922. 10.1126/science.aav8907

Ren, Y., Yue, H., Li, L., Xu, Y., Wang, Z., Xin, Z., & Lin, T. (2018). Identification and characterization of circRNAs involved in the regulation of low nitrogen-promoted root growth in hexaploid wheat. Biological Research, 51(1), 43. 10.1186/s40659-018-0194-3

Riemer, E., Pullagurla, N. J., Yadav, R., Rana, P., Jessen, H. J., Kamleitner, M., Schaaf, G., & Laha, D. (2022). Regulation of plant biotic interactions and abiotic stress responses by inositol polyphosphates. Frontiers in Plant Science, 13. 10.3389/fpls.2022.944515

Robinson, M. D., McCarthy, D. J., & Smyth, G. K. (2010). edgeR: A Bioconductor package for differential expression analysis of digital gene expression data. Bioinformatics, 26(1), 139–140. 10.1093/bioinformatics/btp616

Salmena, L., Poliseno, L., Tay, Y., Kats, L., & Pandolfi, P. P. (2011). A ceRNA Hypothesis: The Rosetta Stone of a Hidden RNA Language? Cell, 146(3), 353–358. 10.1016/j.cell.2011.07.014

Singh, A., Singh, N. B., Yadav, V., Bano, C., Niharika, Khare, S., & Yadav, R. K. (2021). Nod factor signaling in legume-*Rhizobium* symbiosis: Specificity and molecular genetics of nod factor signaling. In V. P. Singh, S. Singh, D. K. Tripathi, S. M. Prasad, R. Bhardwaj, & D. K. Chauhan (Eds.), Abiotic Stress and Legumes (pp. 33–67). Academic Press. 10.1016/B978-0-12-815355-0.00003-5

Singh, S., Katzer, K., Lambert, J., Cerri, M., & Parniske, M. (2014). CYCLOPS, A DNA-Binding Transcriptional Activator, Orchestrates Symbiotic Root Nodule Development. Cell Host & Microbe, 15(2), 139–152. 10.1016/j.chom.2014.01.011

Soyano, T., Hirakawa, H., Sato, S., Hayashi, M., & Kawaguchi, M. (2014). NODULE INCEPTION creates a long-distance negative feedback loop involved in homeostatic regulation of nodule organ production. Proceedings of the National Academy of Sciences of the United States of America, 111(40), 14607–14612. 10.1073/pnas.1412716111

Stracke, S., Kistner, C., Yoshida, S., Mulder, L., Sato, S., Kaneko, T., Tabata, S., Sandal, N., Stougaard, J., Szczyglowski, K., & Parniske, M. (2002). A plant receptor-like kinase required for both bacterial and fungal symbiosis. Nature, 417(6892), 959–962. 10.1038/nature00841

Sullivan, J. T., Brown, S. D., Yocum, R. R., & Ronson, C. W. (2001). The bio operon on the acquired symbiosis island of Mesorhizobium sp. Strain R7A includes a novel gene involved in pimeloyl-CoA synthesisThe GenBank accession number for the sequence reported in this paper is AF311738. Microbiology, 147(5), 1315–1322. 10.1099/00221287-147-5-1315

Suzaki, T., Ito, M., Yoro, E., Sato, S., Hirakawa, H., Takeda, N., & Kawaguchi, M. (2014). Endoreduplication-mediated initiation of symbiotic organ development in Lotus japonicus. Development, 141(12), 2441–2445. 10.1242/dev.107946

Tatsukami, Y., Nambu, M., Morisaka, H., Kuroda, K., & Ueda, M. (2013). Disclosure of the differences of Mesorhizobium loti under the free-living and symbiotic conditions by comparative proteome analysis without bacteroid isolation. BMC Microbiology, 13(1), 180. 10.1186/1471-2180-13-180

Thomas, L., & Rahman, Z. (2020). A Genome-Wide Investigation on Symbiotic Nitrogen-Fixing Bacteria in Leguminous Plants. In A. Varma, S. Tripathi, & R. Prasad (Eds.), Plant Microbe Symbiosis (pp. 55–73). Springer International Publishing. 10.1007/978-3-030-36248-5_4

Tirichine, L., Imaizumi-Anraku, H., Yoshida, S., Murakami, Y., Madsen, L. H., Miwa, H., Nakagawa, T., Sandal, N., Albrektsen, A. S., Kawaguchi, M., Downie, A., Sato, S., Tabata, S., Kouchi, H., Parniske, M., Kawasaki, S., & Stougaard, J. (2006). Deregulation of a Ca2+/calmodulin-dependent kinase leads to spontaneous nodule development. Nature, 441(7097), 1153–1156. 10.1038/nature04862

Tsikou, D., Yan, Z., Holt, D. B., Abel, N. B., Reid, D. E., Madsen, L. H., Bhasin, H., Sexauer, M., Stougaard, J., & Markmann, K. (2018). Systemic control of legume susceptibility to rhizobial infection by a mobile microRNA. Science, 236(October), 233–236. 10.1126/science.aat6907(2018)

Utley, D., Edwards, B., Budnick, A., Grotewold, E., & Sederoff, H. (2025). Camelina circRNA landscape: Implications for gene regulation and fatty acid metabolism. The Plant Genome, 18(1), e20537. 10.1002/tpg2.20537

Vromman, M., Anckaert, J., Bortoluzzi, S., Buratin, A., Chen, C.-Y., Chu, Q., Chuang, T.-J., Dehghannasiri, R., Dieterich, C., Dong, X., Flicek, P., Gaffo, E., Gu, W., He, C., Hoffmann, S., Izuogu, O., Jackson, M. S., Jakobi, T., Lai, E. C.,… Volders, P.-J. (2023). Large-scale benchmarking of circRNA detection tools reveals large differences in sensitivity but not in precision. Nature Methods. 10.1038/s41592-023-01944-6

Wang, Y., Yang, Y., Jiang, X., Shi, J., Yang, Y., Jiang, S., Li, Z., Wang, D., & Chen, Z. (2022). The Sequence and Integrated Analysis of Competing Endogenous RNAs Originating from Tea Leaves Infected by the Pathogen of Tea Leaf Spot, Didymella segeticola. Plant Disease, 106(4), 1286–1290. 10.1094/PDIS-06-21-1324-A

Wu, Z., Huang, W., Qin, E., Liu, S., Liu, H., Grennan, A. K., Liu, H., & Qin, R. (2020). Comprehensive Identification and Expression Profiling of Circular RNAs During Nodule Development in Phaseolus vulgaris. Frontiers in Plant Science, 11. 10.3389/fpls.2020.587185

Xia, K., Pan, X., Zeng, X., & Zhang, M. (2023). Xoo-responsive transcriptome reveals the role of the circular RNA133 in disease resistance by regulating expression of OsARAB in rice. Phytopathology Research, 5(1), 33. 10.1186/s42483-023-00188-8

Xu, C., Li, J., Wang, H., Liu, H., Yu, Z., & Zhao, Z. (2023). Whole-Transcriptome Sequencing Reveals a ceRNA Regulatory Network Associated with the Process of Periodic Albinism under Low Temperature in Baiye No. 1 (Camellia sinensis). International Journal of Molecular Sciences, 24(8), Article 8. 10.3390/ijms24087162

Xue, L., Cui, H., Buer, B., Vijayakumar, V., Delaux, P.-M., Junkermann, S., & Bucher, M. (2015). Network of GRAS Transcription Factors Involved in the Control of Arbuscule Development in Lotus japonicus. Plant Physiology, 167(3), 854–871. 10.1104/pp.114.255430

Yano, K., Yoshida, S., Müller, J., Singh, S., Banba, M., Vickers, K., Markmann, K., White, C., Schuller, B., Sato, S., Asamizu, E., Tabata, S., Murooka, Y., Perry, J., Wang, T. L., Kawaguchi, M., Imaizumi-Anraku, H., Hayashi, M., & Parniske, M. (2008). CYCLOPS, a mediator of symbiotic intracellular accommodation. Proceedings of the National Academy of Sciences, 105(51), 20540–20545. 10.1073/pnas.0806858105

Zeng, X., Lin, W., Guo, M., & Zou, Q. (2017). A comprehensive overview and evaluation of circular RNA detection tools. PLoS Computational Biology, 13(6), e1005420. 10.1371/journal.pcbi.1005420

Zeng, Z., Liu, Y., Feng, X.-Y., Li, S.-X., Jiang, X.-M., Chen, J.-Q., & Shao, Z.-Q. (2023). The RNAome landscape of tomato during arbuscular mycorrhizal symbiosis reveals an evolving RNA layer symbiotic regulatory network. Plant Communications, 4(1). 10.1016/j.xplc.2022.100429

Zhang, D., Ma, Y., Naz, M., Ahmed, N., Zhang, L., Zhou, J.-J., Yang, D., & Chen, Z. (2024). Advances in CircRNAs in the Past Decade: Review of CircRNAs Biogenesis, Regulatory Mechanisms, and Functions in Plants. Genes, 15(7), Article 7. 10.3390/genes15070958

Zheng, Y., & Zhao, F. (2018). Detection and Reconstruction of Circular RNAs from Transcriptomic Data. In C. Dieterich & A. Papantonis (Eds.), Circular RNAs: Methods and Protocols (pp. 1–8). Springer. 10.1007/978-1-4939-7562-4_1

Zhou, F., Liu, Y., Wang, W., Wu, L., Ma, J., Zhang, S., Wang, J., Feng, F., Yuan, H., & Huang, X. (2022). Identification and functional prediction of CircRNAs of developing seeds in high oleic acid sunflower (Helianthus annuus L.). Acta Physiologiae Plantarum, 45(1), 13. 10.1007/s11738-022-03482-8

Zhu, C., Yuan, T., Yang, K., Liu, Y., Li, Y., & Gao, Z. (2023). Identification and characterization of CircRNA-associated CeRNA networks in moso bamboo under nitrogen stress. BMC Plant Biology, 23(1), 142. 10.1186/s12870-023-04155-5

