## Supplemental Figures for "Circular RNAs in Lotus japonicus Responses to Nutrient Supply and Symbiotic Interactions"

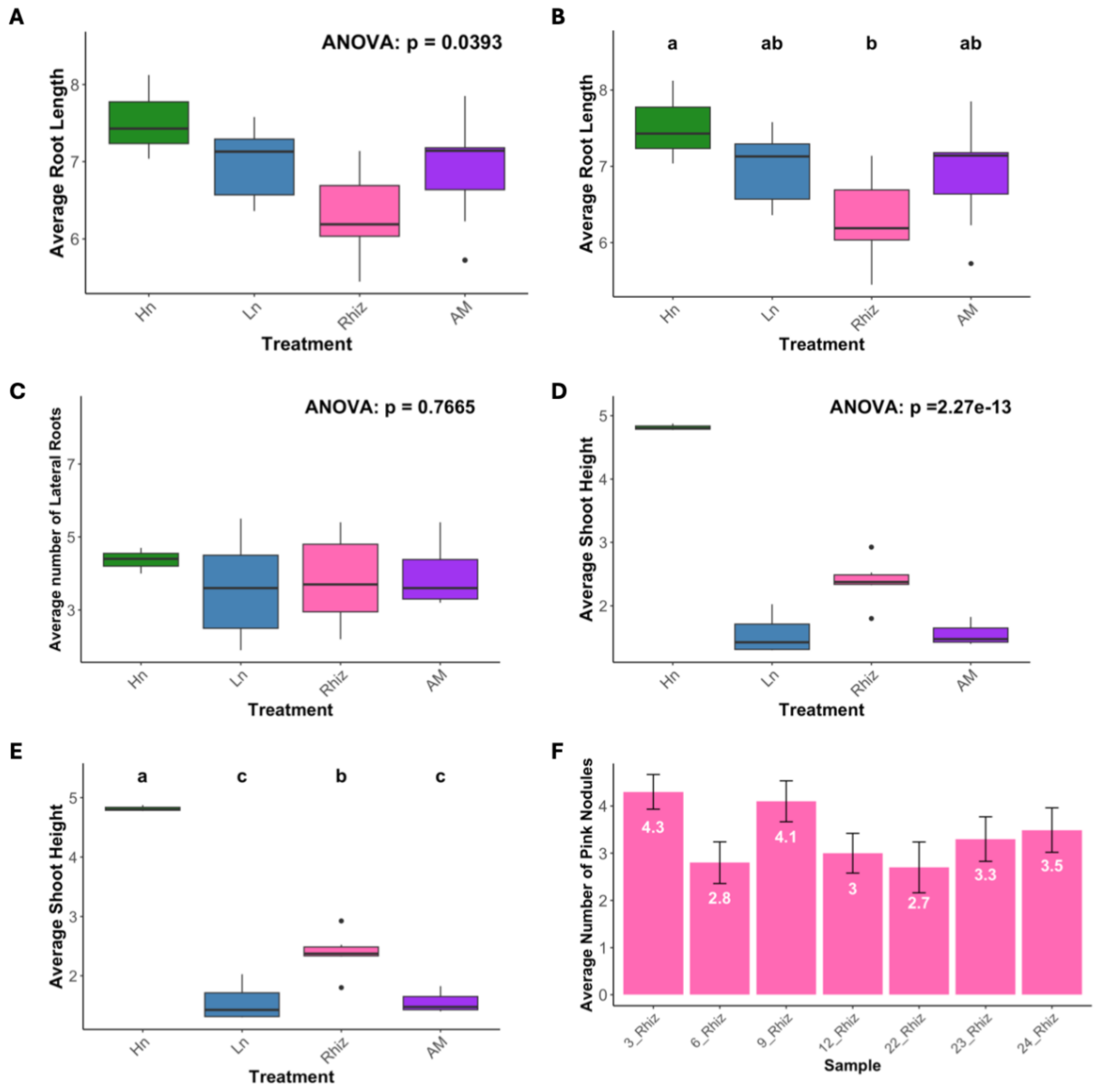


**Supplemental Figure 1**. **Phenotyping of root length, lateral roots, shoot height, and nodule count.** ANOVA (A) and Tukey HSD (Honestly Significant Difference) of root length (B). ANOVA results of lateral roots (C). ANOVA (D) and Tukey HSD (E) of shoot height. Average number of nodules per bioreplicate for the Rhiz treatment.

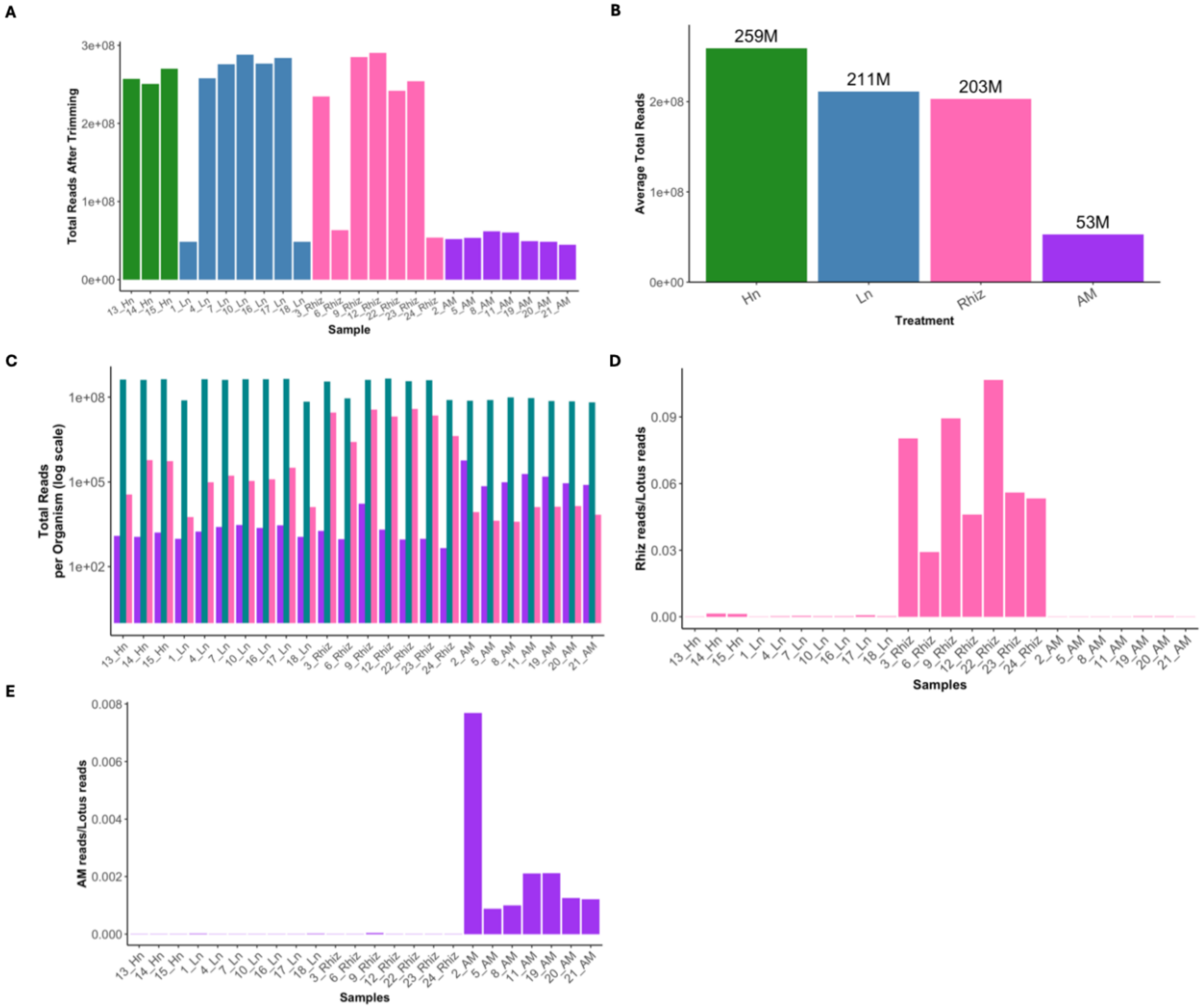


**Supplemental Figure 2. Assigned reads.** (A) Total number of reads per sample after trimming, illustrating the distribution of sequencing depth across all samples. (B) Average number of reads per sample, providing an overview of the sequencing consistency across treatments. Note that samples associated with AM were subjected to shallow sequencing (40M reads per sample) only, resulting in comparatively lower read counts. (C) Log-transformed total reads per sample from Bbsplit analysis, showing the distribution of assigned reads for *L. japonicus*, *M.loti*, and AM, illustrating the relative contribution of each organism to the total sequencing output (D) Ratio of organism-specific reads for *M.loti* and (E) *R. iregularis*.

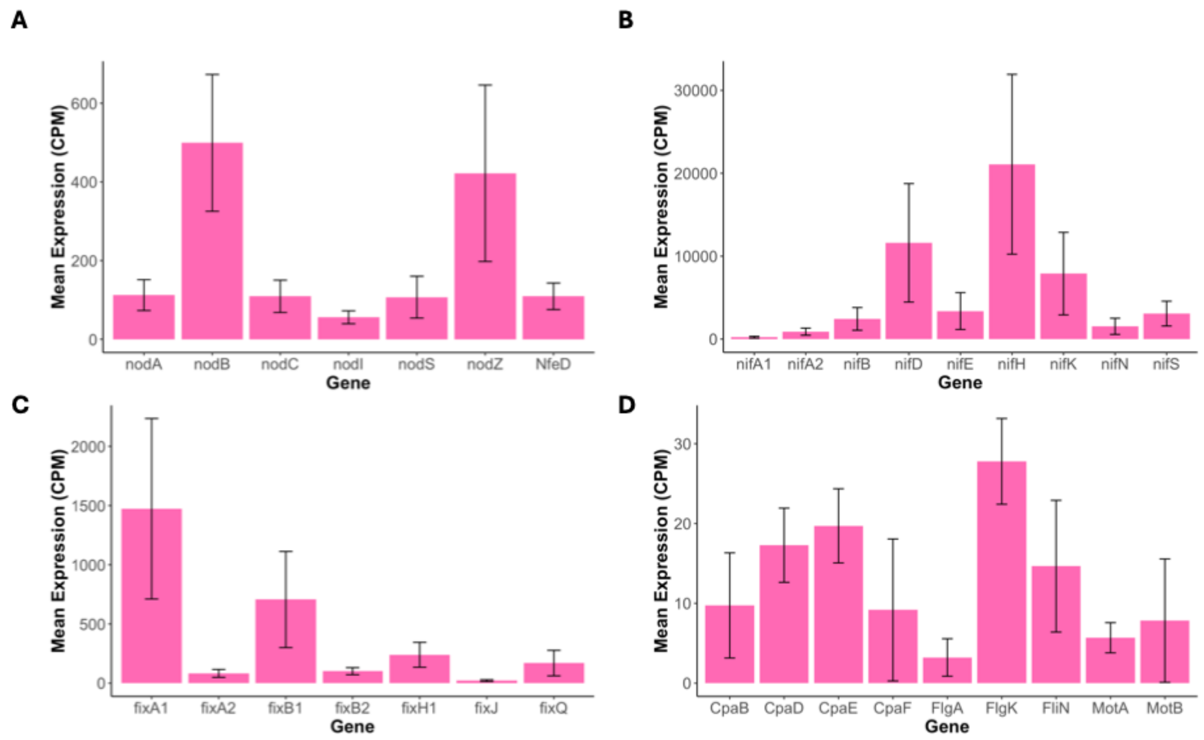


**Supplemental Figure 3. Mean Expression (CPM) of nod, fix, nif, and structural flagellar and pili transcripts in *M. loti* as sampled from the roots of *L. japonicus*.** Nod factor genes (A) and genes encoding flagellar and pili structural elements (D) are associated mainly or exclusively with free-living bacteria, while the nitrogen fixation genes (nif and fix, B and C), present in bacteroids, are also highly expressed. Supplemental File 5 contains raw expression data from the *M. loti* transcriptome.

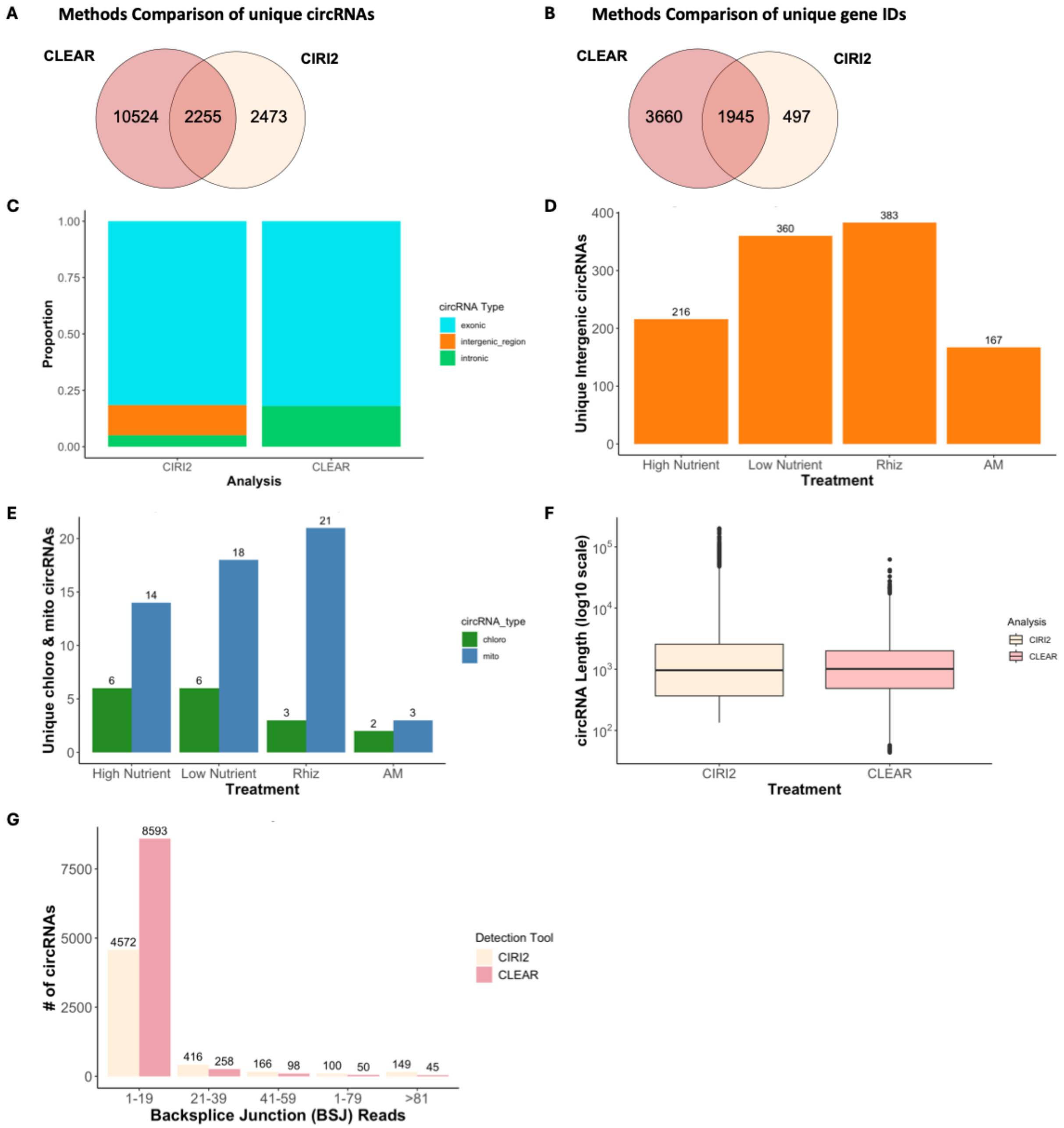


**Figure 4. CircRNA landscape. (**A) Comparison of unique nuclear circRNAs from the two detection tools, CircExplorer3 (CLEAR) and CIRI2. (B) Comparison of unique cognate gene IDs from CLEAR and CIRI2. (C) Proportion of the three types of circRNAs detected from the two detection tools, nuclear circRNAs only. (D) Total number of unique intergenic circRNAs in each treatment. (E) Total number of unique chloroplast and mitochondria circRNAs from each treatment. (F) Box and Whisker plot of circRNA length between CIRI2 and CLEAR. (G) Number of CLEAR and CIRI2 unique circRNAs and the number of BSJ reads.

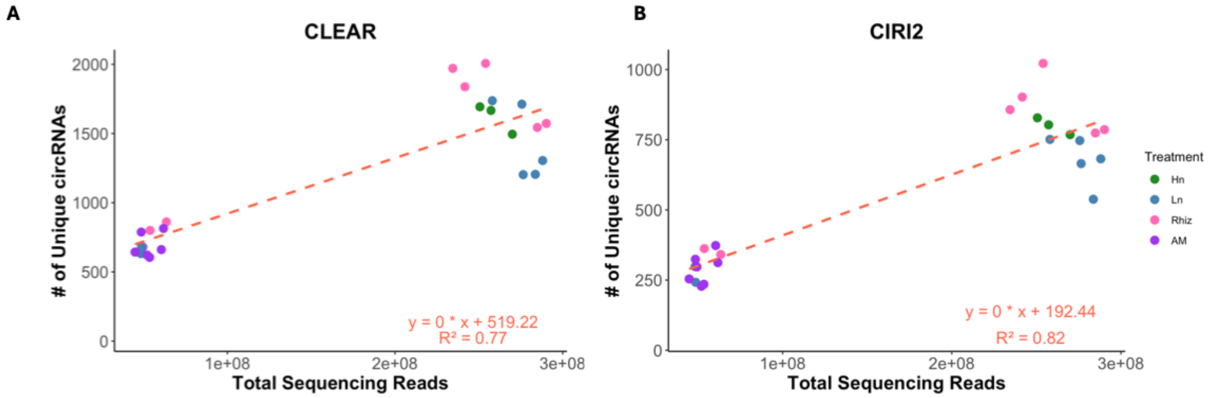


**Supplemental Figure 5. Comparison of read depth and circRNAs** per sample as identified by either CLEAR (A) or CIRI2 (B).

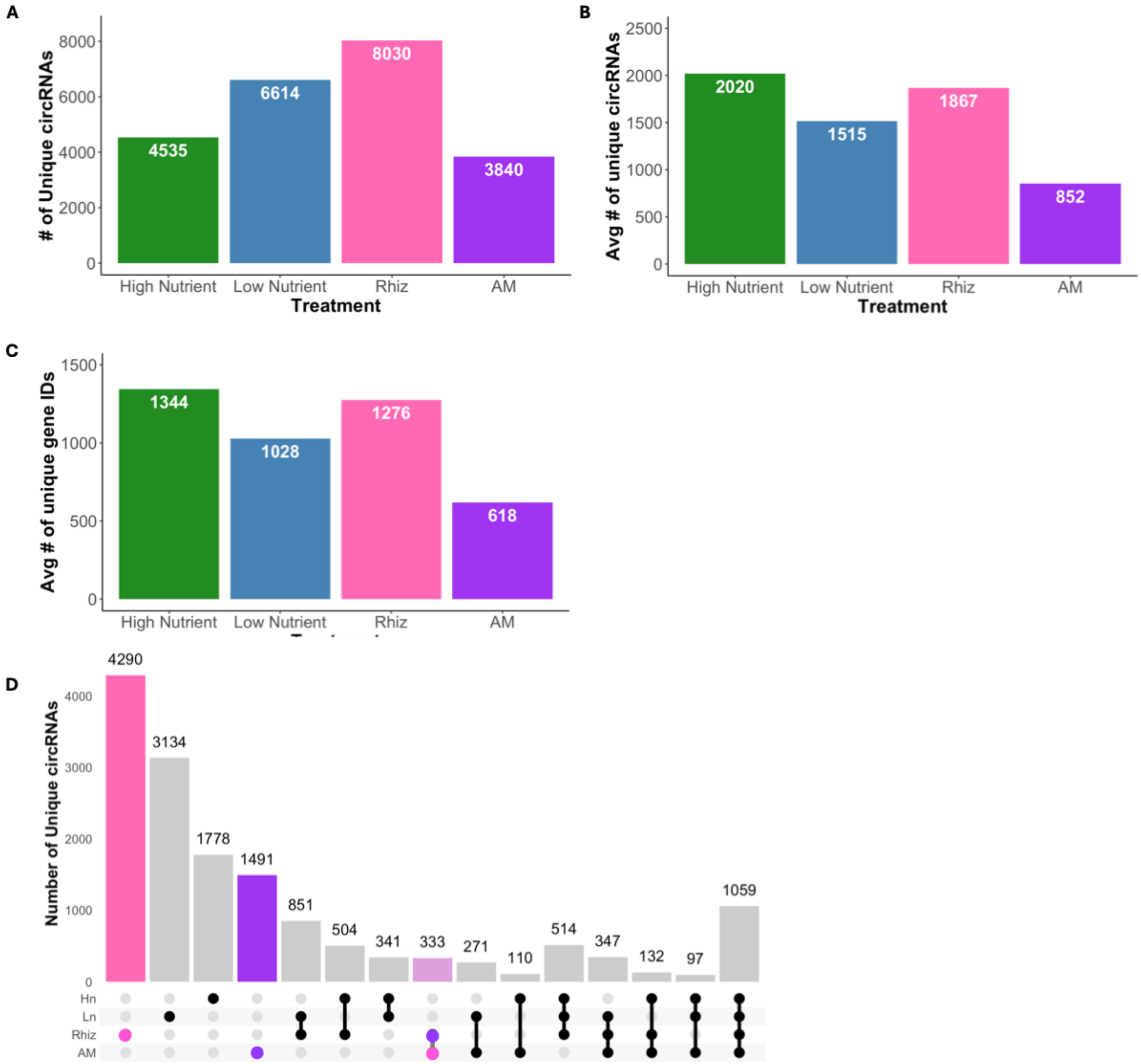


**Supplemental Figure 6. The qualitative and quantitative abundance of circRNAs and their cognate gene IDs.** (A) Number of unique circRNAs per treatment. (B) Average number of unique circRNAs per treatment. (C) Average number of genes generating circRNA per treatment. (D) Distribution of unique circRNAs across treatments. Upset plot showing the number of unique circRNAs for specific treatments and combinations of treatments. Rhiz (pink), AM (purple), combination of the two (plum).

**Supplemental Table 1. Select Treatment Specific circRNA.** Conserved (≥ 4 replicates in Ln, Rhiz, and AM, or ≥ 2 replicates in Hn) circRNAs from a treatment were compared to all circRNAs of the other treatments (Figure 3). Select treatment-specific circRNAs are listed in this table. Complete table found in Supplemental File 7.

| Treatment | circRNA | Lj gene ID | Functional annotation |
|---|---|---|---|
| **cons_Hn** | LjG1.1_chr1:129532249-129532599 | LotjaGi1g1v0724100 | 30S ribosomal protein S10 |
| **cons_Hn, all Ln** | LjG1.1_chr1:105469038-105471214 | LotjaGi1g1v0527800 | Argininosuccinate synthase |
| | LjG1.1_chr1:18680735-18681182 | LotjaGi1g1v0118200 | Asparagine synthetase |
| | LjG1.1_chr3:58252953-58253127, LjG1.1_chr3:58252954-58253127 | LotjaGi3g1v0288100 | Xanthine dehydrogenase |
| **cons_Ln, all Hn** | LjG1.1_chr6:60727432-60728319, LjG1.1_chr6:60727433-60728319 | LotjaGi6g1v0290100 | Leucine-rich repeat receptor-like protein kinase |
| **cons_Ln, all Rhiz** | LjG1.1_chr2:94856012-94856162 | LotjaGi2g1v0455500_LC | Acyl-CoA N-acyltransferase with RING/FYVE/PHD-type zinc finger protein |
| **cons_Rhiz** | LjG1.1_chr1:104035513-104036488,LjG1.1_chr1:104035514-104036488 | LotjaGi1g1v0517300 | 2-oxoglutarate (2OG) and Fe(II)-dependent oxygenase superfamily protein |
| | LjG1.1_chr3:91852722-91853514 | LotjaGi3g1v0500200 | RNA-binding protein |
| | LjG1.1_chr4:81678626-81680400, LjG1.1_chr4:81678627-81680400, LjG1.1_chr4:81678627-81680396 | LotjaGi4g1v0443300 | ATP-dependent 6-phosphofructokinase |
| | LjG1.1_chr5:43488209-43488486, LjG1.1_chr5:43488210-43488486 | LotjaGi5g1v0169900 | Type I inositol-1,4,5-trisphosphate 5-phosphatase CVP2 |
| | LjG1.1_chr5:58943587-58947147, LjG1.1_chr5:58943588-58947147 | LotjaGi5g1v0275400 | Aminodeoxychorismate synthase |
| **cons_Rhiz, all Hn** | LjG1.1_chr6:36811083-36811521 | LotjaGi6g1v0130800 | Asparagine synthetase |
| **cons_Rhiz, all Ln** | LjG1.1_chr1:121699751-121700534, LjG1.1_chr1:121699752-121700534 | LotjaGi1g1v0657000 | Transcription elongation factor SPT6 |
| | LjG1.1_chr1:51353902-51354488, LjG1.1_chr1:51353903-51354488 | LotjaGi1g1v0224700 | Glutamate synthase, putative |
| | LjG1.1_chr4:25449377-25450306, LjG1.1_chr4:25449378-25450306 | LotjaGi4g1v0153200 | Uridine kinase |
| | LjG1.1_chr4:65590834-65591669, LjG1.1_chr4:65590835-65591669 | LotjaGi4g1v0283700 | Dicer-like 3 |
| **cons_Rhiz, all AM** | LjG1.1_chr2:80616495-80617260 | LotjaGi2g1v0330500 | Receptor kinase; SYMRK |

| | | | |
|---|---|---|---|
| **cons AM, all Rhiz** | LjG1.1_chr2:79333642-79336472, LjG1.1_chr2:79333643-79336472 | LotjaGi2g1v0318400 | AP-1 complex subunit gamma-2 |

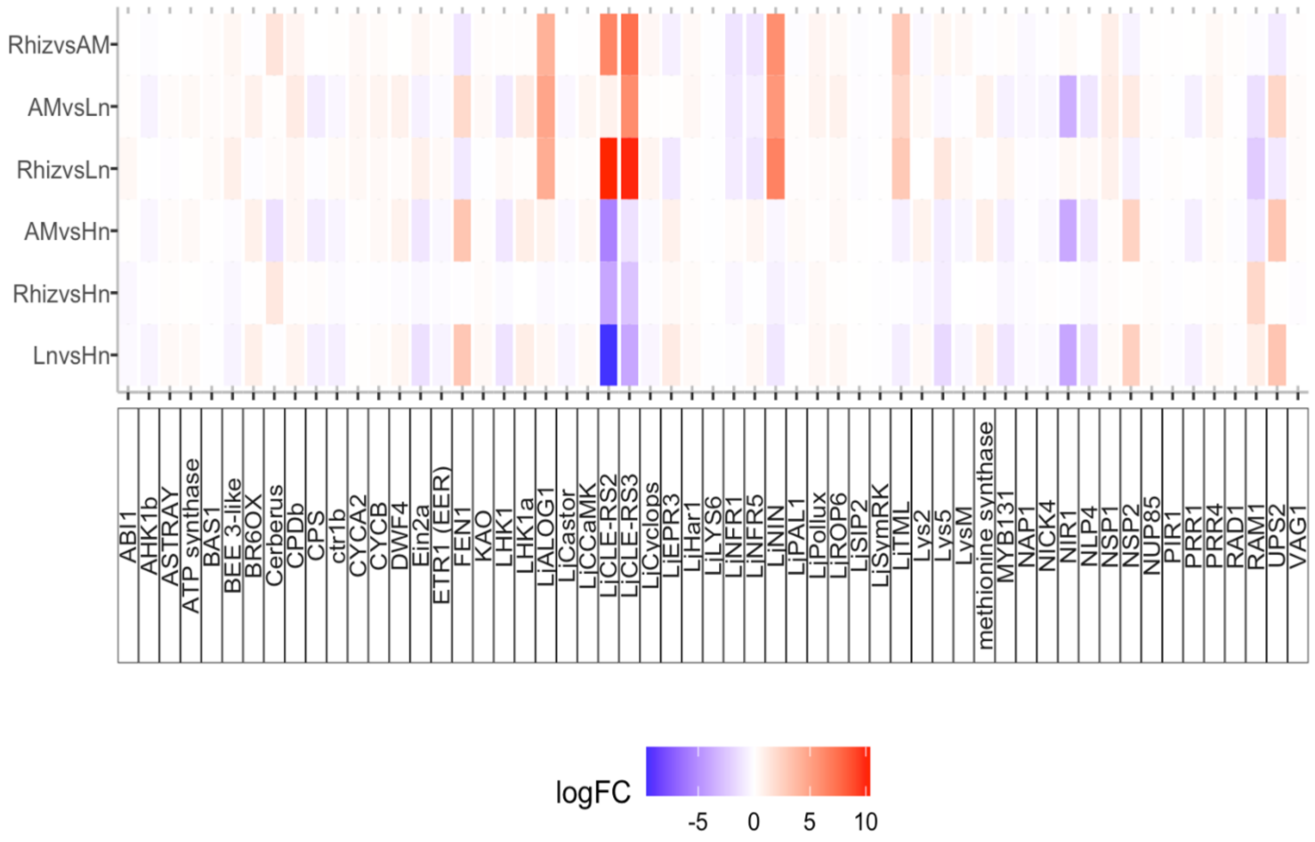


**Supplemental Figure 7. Heatmap of the differential expression of CSP genes in different comparisons.** Color represents the logFC of CSP genes (x-axis) in the different comparisons (y-axis). No FDR cutoff applied

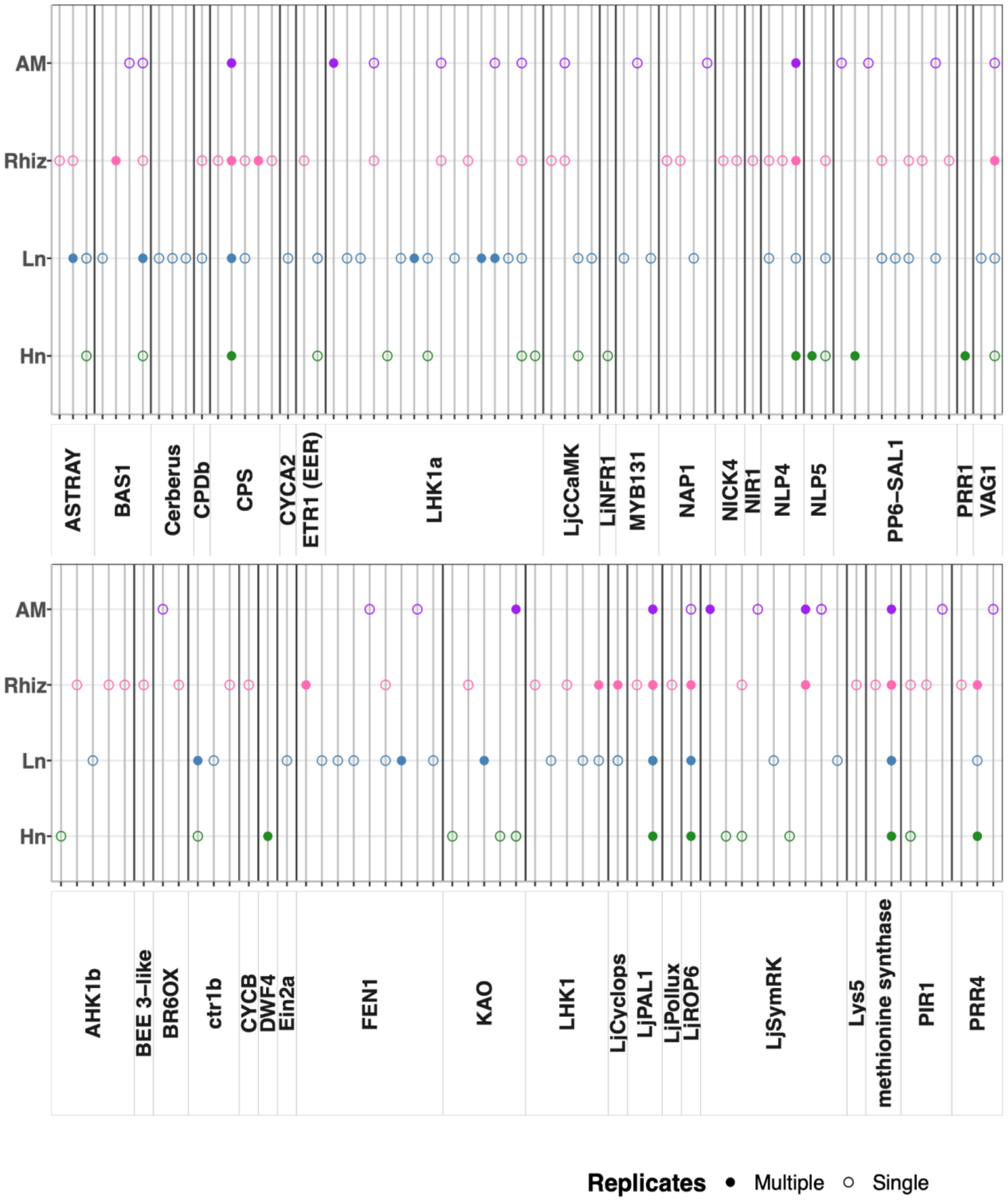


**Supplemental Figure 8. Common Symbiosis Pathway genes and associated circRNAs.** 38 CSP genes with their associated circRNAs. Light grey lines represent different circRNAs associated with a particular gene. Circles on the light grey horizontal grid lines represent how many replicates the circRNAs were present in. All data in Supplemental File 6.

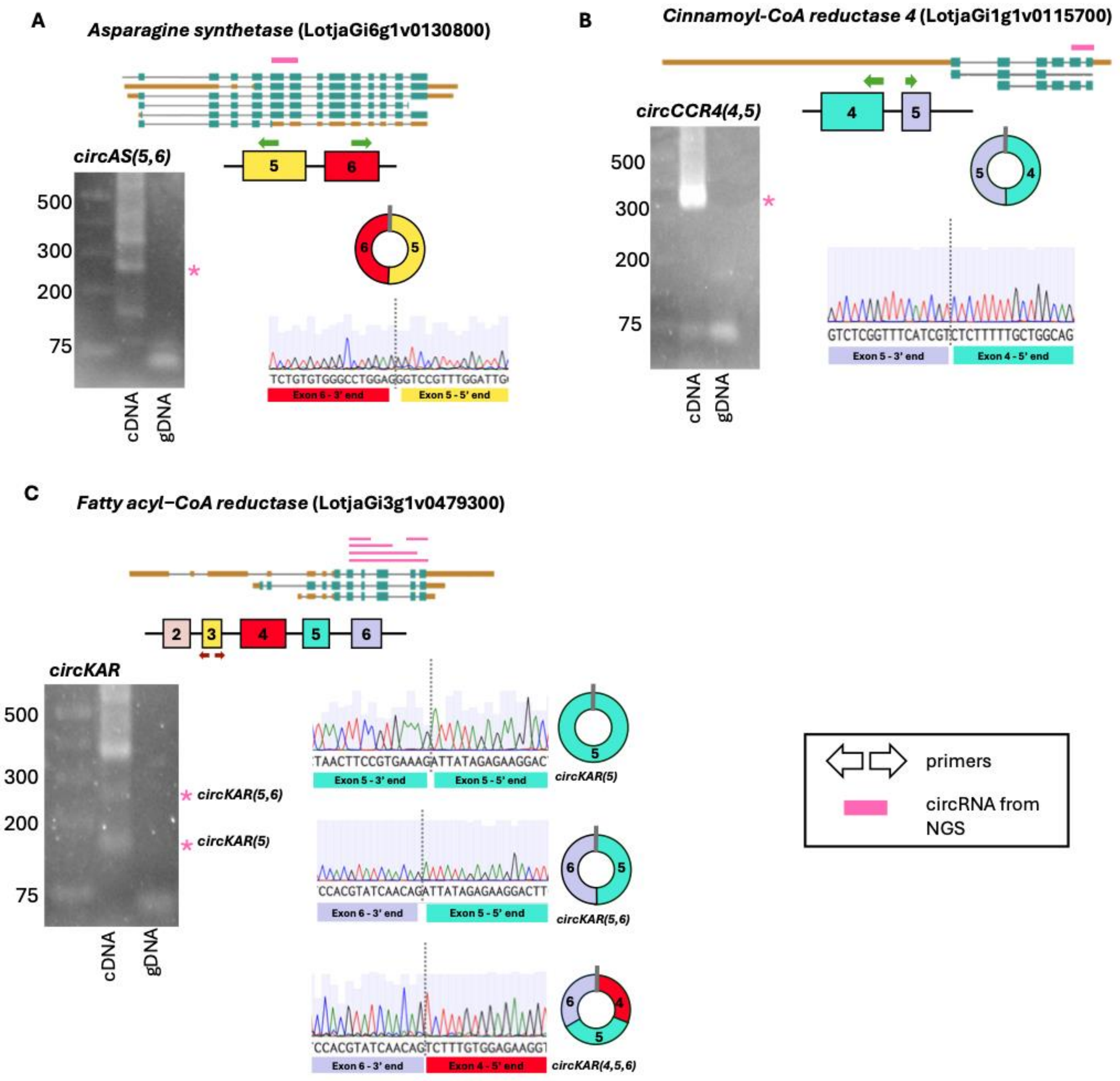


**Supplemental Figure 9. Confirmation of select circRNAs.** CircRNAs were validated (A) using divergent primers designed to capture the backsplice junction (orange arrows) and to maximize the potential of capturing multiple circs that may have a shared exon (red arrows). (B-D) Examples of validated circRNA using RT-PCR and Sanger sequencing. Pink bars above gene schematic represent circRNAs identified from NGS. Arrows represent the location of primers (not drawn to scale). Pink asterisk represents the band in the cDNA lane that was Sanger sequenced. Gray dotted/solid line represents the backsplice junction. Full image of gel (Supplemental Figure 11). All coordinates of validated circRNAs can be found in Supplemental File 2.

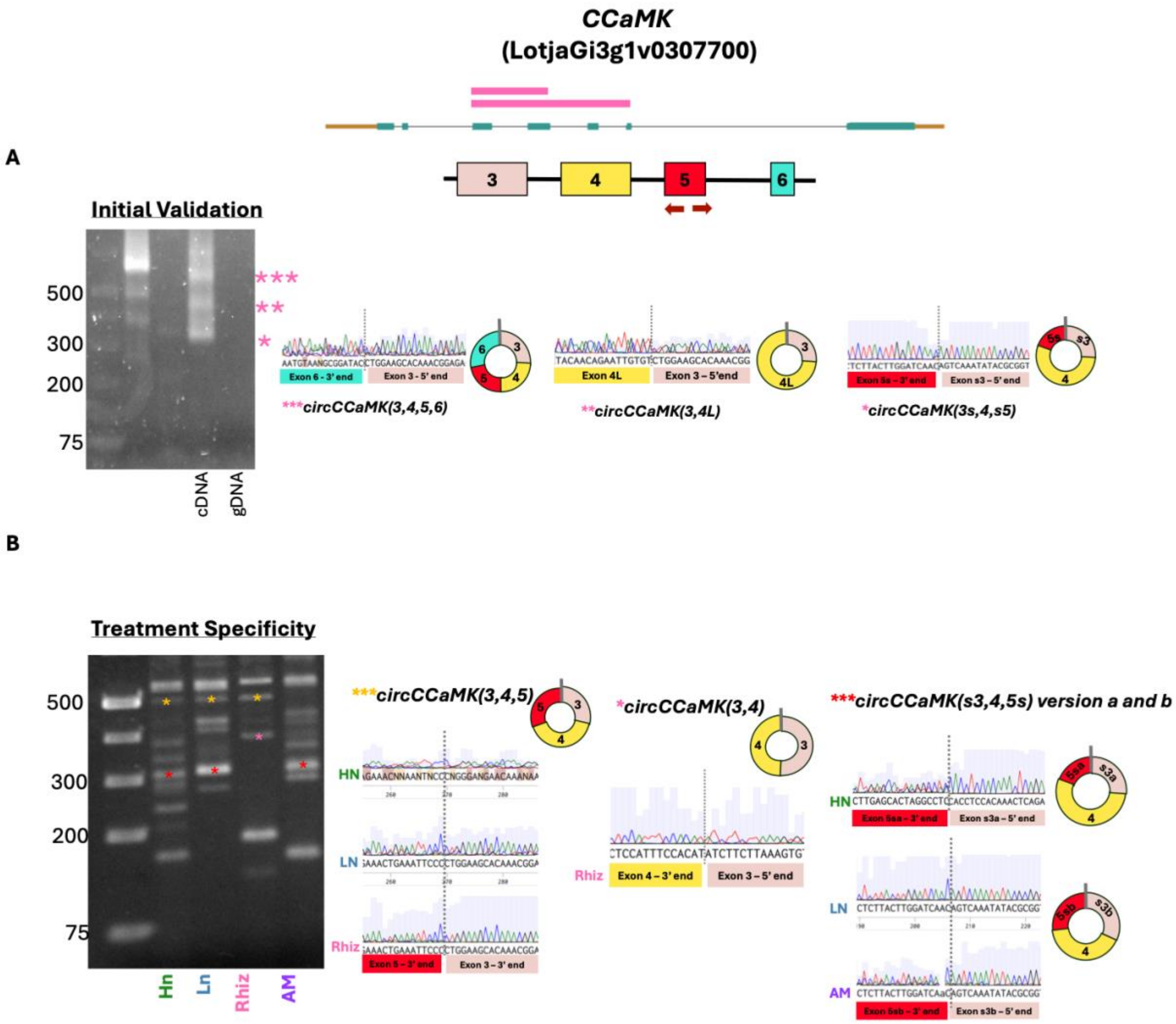


**Supplemental Figure 10. Validation of *CCamK* circRNAs.** (A) BSJ sequences from the initial 3 validated circRNAs using Rhiz tissue. (B) BSJ sequences for all the treatment-specific circRNAs. All circRNAs shown here correspond to those listed in the tables of Figure 5C.

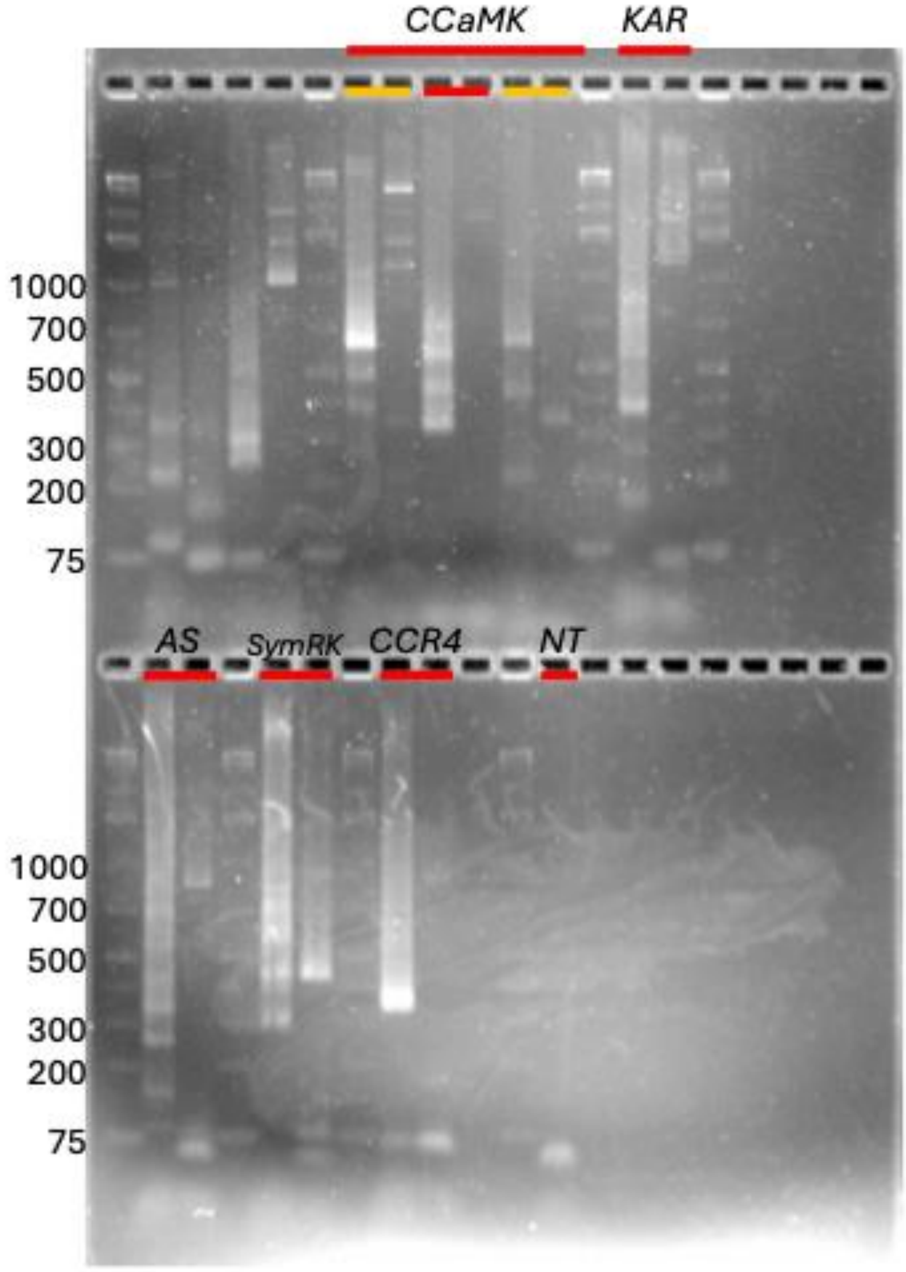


**Supplemental Figure 11. Full image of gel electrophoresis corresponding to Supplemental Figures 9 and 10A.** Red lines over lanes represent lanes in which cDNA (left lane) and gDNA(right lane) are present for that particular gene. For *CCaMK*, there were three different primer pairs used (indicated by the two gold and one red line under *CCaMK*). The smaller red line represents the primer pair chosen to be highlighted in this study. Gel was prepared as 2% agarose in TBE, run at 200V, and post-stained with GelGreen. The ladder used is Generuler 1kb DNA ladder. Unlabelled lanes do not pertain to the information in this publication. Primers used can be found in Supplemental File 2.

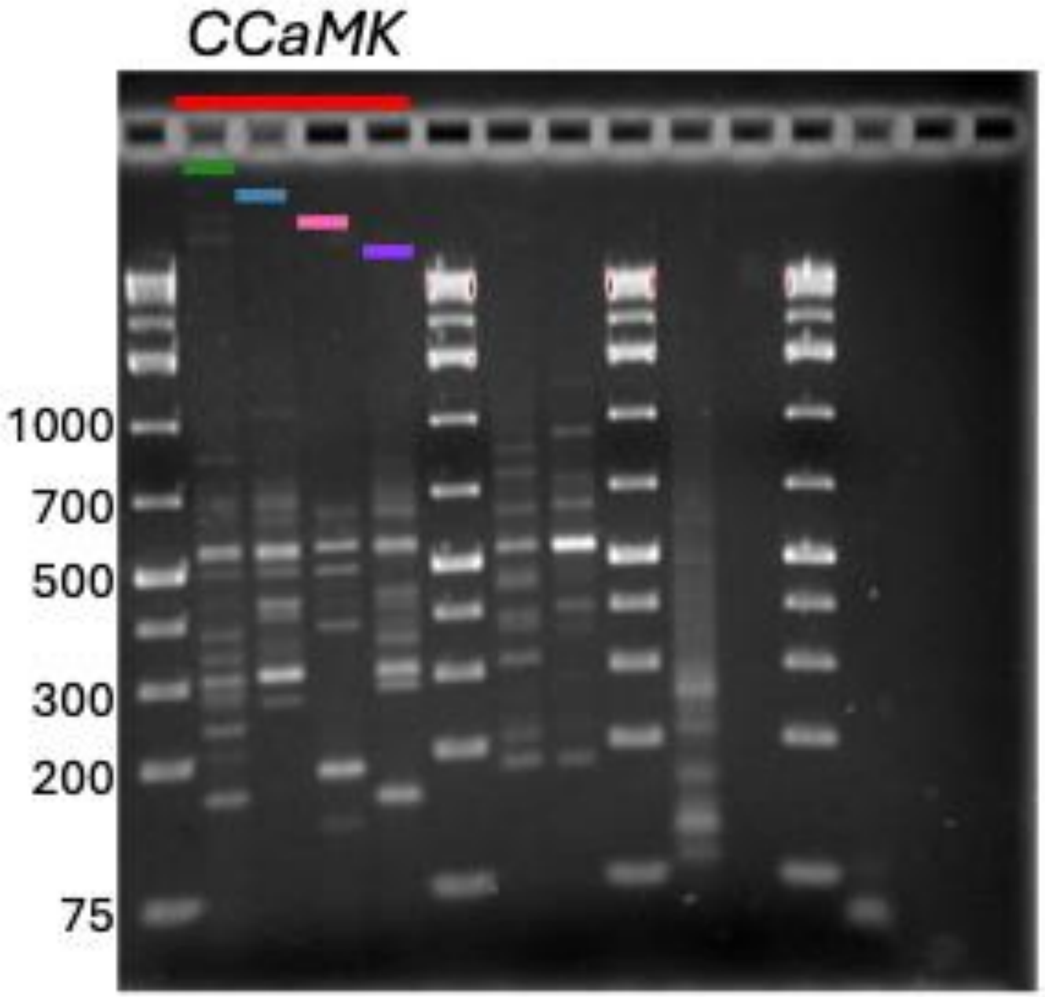


**Supplemental Figure 12. Full image of gel electrophoresis corresponding to Figure 6C.** Four lanes to the left of the gel represent products using the same primer pair from the selected pair in Supplemental Figure 5 (small red line). The small colored lines here represent the cDNA from specific treatments: Green - Hn, blue- Ln, pink-Rhiz, and purple - AM.

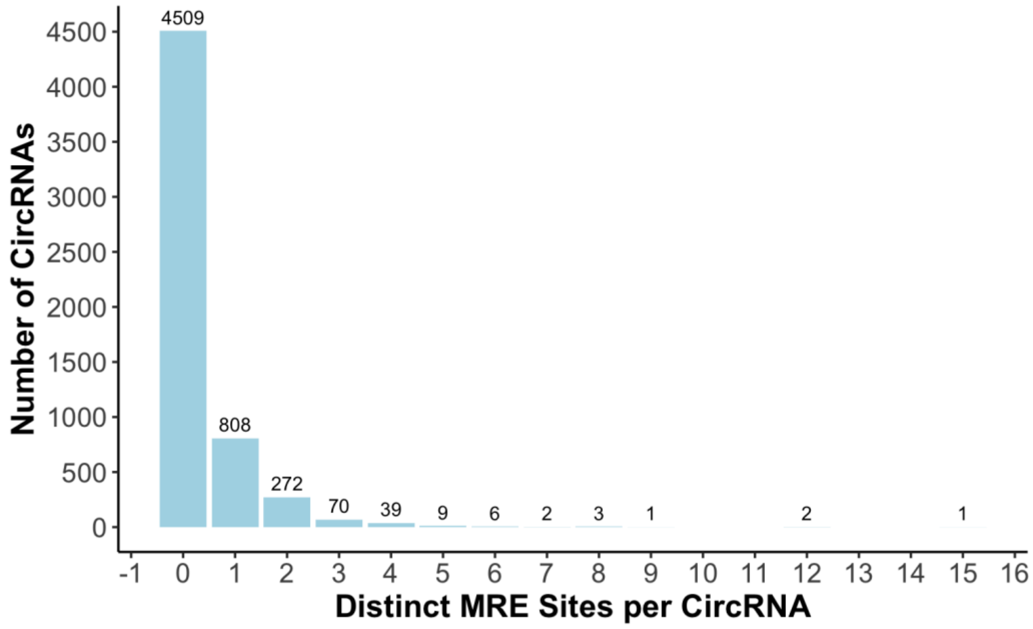


**Supplemental Figure 13. Distribution of circRNAs to unique MREs.**

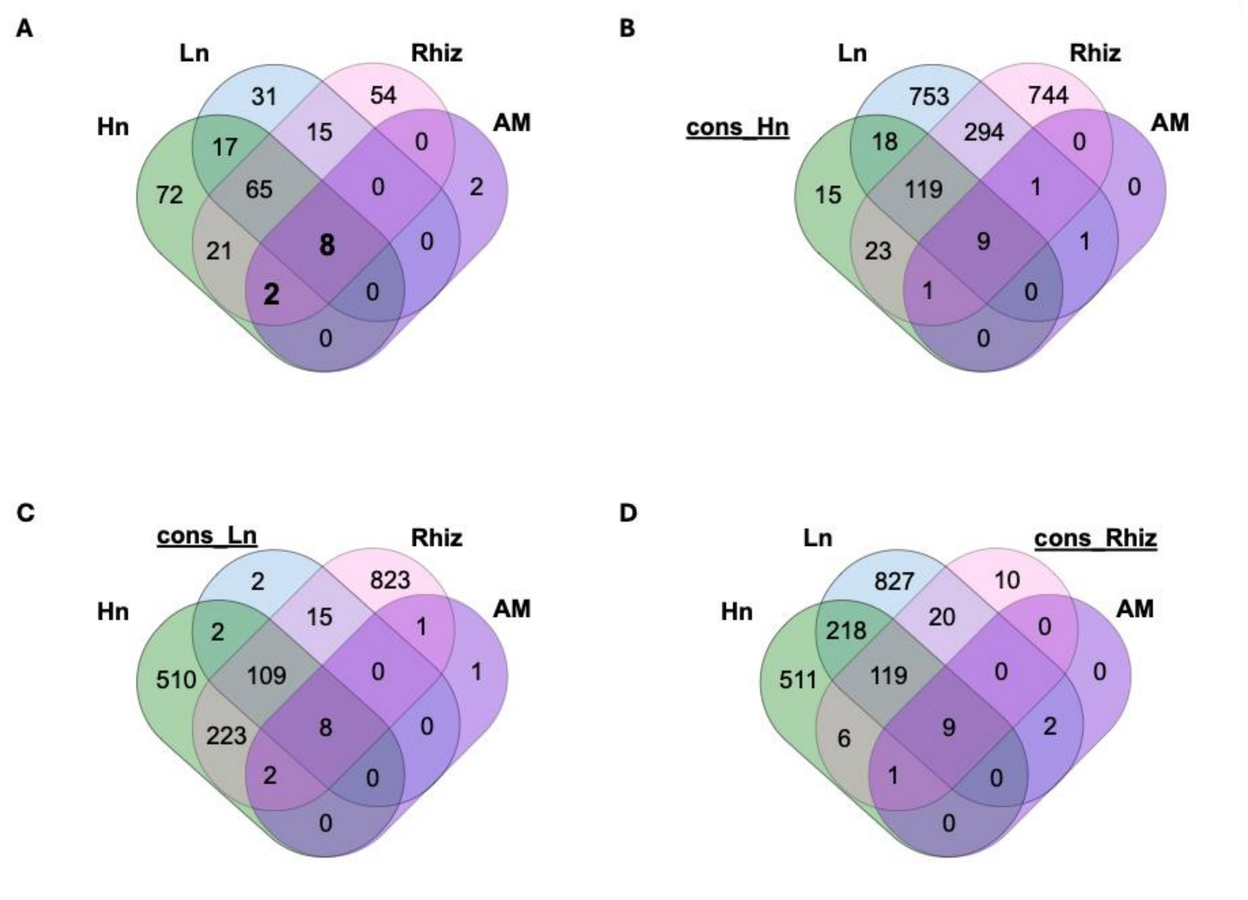


**Supplemental Figure 14. Comparison of circRNAs with MREs.** (A) Comparison of circRNAs with MREs that were present in at least three samples of the Ln and Rhiz treatments, at least two samples of the Hn treatment, and any of the AM treatments. Numbers in bold represent those circRNAs with MREs present in most samples or considered conserved among treatments. Info about circRNAs in bold is in Supplemental Tables 3 and 4.

**Supplemental Table 2. CircRNAs with MREs.** These circRNAs correspond to the eight circRNAs with MREs that are conserved in most treatments and that are shared between AM, Hn, and Rhiz. Corresponding to the numbers in bold in Figure 16A.

| | circRNA | Lj gene ID | Ath homeolog | Functional Annotation |
|---|---|---|---|---|
| **Shared among all (8)** | LjG1.1_chr3:89378043-89379487 | LotjaGi3g1v0477400 | AT1G08320 | transcription factor-like protein; bZIP transcription factor family |
| | LjG1.1_chr2:15791234-15791927 | LotjaGi2g1v0067300 | AT1G23230 | Mediator of RNA polymerase II transcription subunit 23 |
| | LjG1.1_chr4:78251887-78252651 | LotjaGi4g1v0406300 | AT5G64740 | Cellulose synthase;(Nucleotide-diphospho-sugar transferases) |
| | LjG1.1_chr5:1100076-1100474 | LotjaGi5g1v0004500 | AT1G75850 | Vacuolar protein sorting-associated protein 35 |
| | LjG1.1_chr5:2580602-2581319 | LotjaGi5g1v0011300 | AT1G06060 | LisH and RanBPM domains containing protein |
| | LjG1.1_chr1:79803880-79805122 | LotjaGi1g1v0337600 | AT5G50530 | CBS domain-containing protein |
| | LjG1.1_chr3:66004333-66004454 | LotjaGi3g1v0311500_LC.1 | AT4G12540 | V-type ATP synthase alpha chain |
| | LjG1.1_chr1:4953249-4953694 | LotjaGi1g1v0035800.1 | AT2G18950 | Homogentisate phytyltransferase |
| **AM, Hn, Rhiz (2)** | LjG1.1_chr3:90030866-90031579 | LotjaGi3g1v0482400 | AT2G27900 | Coiled-coil domain-containing protein 132 |
| | LjG1.1_chr3:55229888-55231162 | LotjaGi3g1v0278400 | AT1G07705 | CCR4-NOT transcription complex subunit 2 |

**Supplemental Table 3. MRE containing circRNAs that are specific to a treatment.** Several of those treatment-specific circRNAs have more than one miRNA binding site, and a few have miRNA-binding sites generated by the BSJ of the circRNA (red, Offset).

| circRNA | Lj gene ID | Functional Annotation | miRNA_Acc. | miRNA | Inhibition | bsrMRE |
|---|---|---|---|---|---|---|
| **Rhiz-specific** | | | | | | |
| LjG1.1_chr2:48683039-48683266 | LotjaGi2g1v0134400 | Carbamoyl-phosphate synthase large chain | lja-miR11117a-5p | 21 | Cleavage | bsrMRE |
| LjG1.1_chr6:65572369-65572776 | LotjaGi6g1v0342500 | G-type lectin S-receptor-like serine/threonine-protein kinase | lja-miR11161-5p | 24 | Cleavage | ORIG |
| LjG1.1_chr5:63194865-63195172 | LotjaGi5g1v0313400 | Protein SUPPRESSOR OF GENE SILENCING 3; RNA recognition motif XS domain | lja-miR7535 | 19 | Cleavage | ORIG |
| LjG1.1_chr6:64839118-64839252 | LotjaGi6g1v0333700 | Acetolactate synthase | lja-miR11156-5p | 24 | Cleavage | bsrMRE |
| LjG1.1_chr4:23725390-23734714 | LotjaGi4g1v0146800 | Ubiquitin-conjugating enzyme E2 | lja-miR7537 | 22 | Cleavage | ORIG |
| | | | lja-miR11075b-3p | 20 | Translation | ORIG |
| LjG1.1_chr2:88255366-88256235 | LotjaGi2g1v0386800 | ATP-dependent zinc metalloprotease FtsH; Sec14p-like phosphatidylinositol transfer family protein | lja-miR11082l-3p | 21 | Cleavage | ORIG |
| | | | lja-miR11072j-5p | 21 | Cleavage | ORIG |
| | | | lja-miR164-5p | 20 | Cleavage | Orig |
| LjG1.1_chr4:35844601-35846831 | LotjaGi4g1v0176400 | MEDTR Cyclin-dependent kinase | lja-miR11086-5p | 21 | Cleavage | ORIG |

**Table 3.** (continued).

| | | | | | | |
|---|---|---|---|---|---|---|
| LjG1.1_chr5:50442227-50443478 | LotjaGi5g1v0203900 | Protein kinase (Armadillo-type fold) | lja-miR11079-3p | 22 | Cleavage | ORIG |
| LjG1.1_chr3:20475718-20476178 | LotjaGi3g1v0152200 | MACPF domain protein (Membrane attack complex component/perforin (MACPF) domain | lja-miR11106-3p | 21 | Cleavage | bsrMRE |
| | | | lja-miR11072b-3p | 20 | Cleavage | ORIG |
| LjG1.1_chr4:82573774-82575480 | LotjaGi4g1v0453800 | Zinc finger protein, putative (Haemerythrin-likeBTS_ARATH Zinc finger protein BRUTUS);A0ZT53_LOTJA Putative E3 ubiquitin ligase | lja-miR7524 | 21 | Cleavage | ORIG |
| | | | lja-miR7540a | 20 | Cleavage | ORIG |
| | | | lja-miR7540b | 20 | Cleavage | ORIG |
| **Ln specific** | | | | | | |
| LjG1.1_chr3:21235938-21236160 | LotjaGi3g1v0155500 | Transmembrane protein | lja-miR11108q-3p | 24 | Cleavage | ORIG |
| | | | lja-miR477-5p | 22 | Cleavage | ORIG |
| LjG1.1_chr3:92579092-92579284 | LotjaGi3g1v0508600.1 | 3-ketoacyl-CoA synthase | lja-miR11149a-3p | 23 | Cleavage | bsrMRE |
| **Hn specific** | | | | | | |
| LjG1.1_chr5:64989061-64989276 | LotjaGi5g1v0332700 | ATP synthase subunit beta | lja-miR11096a-3p | 21 | Cleavage | ORIG |
| | | | lja-miR11096b-3p | 21 | Cleavage | ORIG |
| LjG1.1_chr3:7920582-7924405 | LotjaGi3g1v0072600.2 | Histidine-containing phosphotransfer protein | lja-miR11129-3p | 22 | Translation | ORIG |

**Table 3.** (continued).

| | | | | | | |
|---|---|---|---|---|---|---|
| LjG1.1_chr2:48682616-48683002 | LotjaGi2g1v0134300 | Carbamoyl-phosphate synthase large chain | lja-miR11117a-5p | 21 | Cleavage | ORIG |
| | | | lja-miR11117a-3p | 21 | Cleavage | ORIG |
| | | | lja-miR11117b-3p | 21 | Cleavage | ORIG |
| | | | lja-miR7533a | 21 | Cleavage | ORIG |
| LjG1.1_chr1:107119239-107120003 | LotjaGi1g1v0542300 | 14-3-3-like protein | lja-miR7533b | 21 | Cleavage | ORIG |
| | | | lja-miR11117a-5p | 21 | Cleavage | ORIG |
| LjG1.1_chr3:91164009-91186085 | LotjaGi3g1v0493300 | ABC transporter ATP-binding protein/permease wht-3; ARATH Protein CROWDED NUCLEI 1;Glycerol-3-phosphate dehydrogenase | lja-miR11108s-3p | 24 | Cleavage | ORIG |
| | | | lja-miR11135a-3p | 22 | Cleavage | ORIG |
| | | | lja-miR11135b-3p | 22 | Cleavage | ORIG |
| | | | lja-miR11135c-3p | 22 | Cleavage | ORIG |
| | | | lja-miR11135d-3p | 24 | Cleavage | ORIG |
| LjG1.1_chr1:7392614-7393114 | LotjaGi1g1v0052500 | Proteasome-associated ECM29-like protein | lja-miR11097d-5p | 22 | Translation | ORIG |
| LjG1.1_chr5:67924498-67924887 | LotjaGi5re1v6146300 | repeat | lja-miR11163-5p | 24 | Cleavage | ORIG |

**Table 3.** (continued).

| | | | | | | |
|---|---|---|---|---|---|---|
| LjG1.1_chr1:10193721-10194447 | LotjaGi1g1v0072600 | Nuclease; AT2G40410.2 Ca(2+)-dependent nuclease family protein | lja-miR11138-3p | 23 | Cleavage | ORIG |
| | | | lja-miR7519 | 22 | Cleavage | ORIG |
| LjG1.1_chr6:5042434-5042733 | LotjaGi6g1v0028000_LC | Bifunctional inhibitor/lipid-transfer protein/seed storage 2S albumin superfamily protein; | lja-miR398-3p | 21 | Translation | bsrMRE |
| | | | lja-miR398-3p | 21 | Cleavage | ORIG |
| | | | lja-miR11125-5p | 22 | Cleavage | ORIG |
| LjG1.1_chr3:25087723-25087819 | LotjaGi3g1v0180300 | F-box protein family | lja-miR11170-3p | 24 | Cleavage | ORIG |
| LjG1.1_chr3:45672664-45678108 | LotjaGi3g1v0259000 | ARATH 187-kDa microtubule-associated protein AIR9; | lja-miR11178-3p | 24 | Cleavage | ORIG |
| | | | lja-miR7536a | 19 | Cleavage | ORIG |
| | | | lja-miR7536b | 19 | Cleavage | ORIG |
| LjG1.1_chr3:89378043-89378309 | LotjaGi3g1v0477400 | transcription factor-like protein (Basic-leucine zipper domain; CAJCA TGACG-sequence-specific DNA-binding protein TGA-2.1, TGA9 | lja-miR11155e-5p | 24 | Cleavage | ORIG |

**Table 3.** (continued).

| | | | | | | |
|---|---|---|---|---|---|---|
| LjG1.1_chr1:3500068-3500629 | no gene | non-coding RNA | lja-miR11114-5p | 23 | Cleavage | ORIG |
| | | | lja-miR11116-3p | 21 | Cleavage | ORIG |
| | | | lja-miR11117a-3p | 21 | Cleavage | ORIG |
| | | | lja-miR11117b-3p | 21 | Cleavage | ORIG |
| LjG1.1_chr5:55941061-55941263 | LotjaGi5g1v0251000 | Beta-glucosidase, putative; TAIR: AT1G02850.2 beta glucosidase 11; | lja-miR11066-3p | 19 | Cleavage | ORIG |
| LjG1.1_chr1:100701952-100702075 | LotjaGi1g1v0493500 | ELMO domain-containing protein, putative; TAIR: AT3G60260.1 ELMO/CED-12 family protein | lja-miR11089a-5p | 21 | Cleavage | ORIG |

**Supplemental Table 4. Treatment-specific microRNA response elements (MREs) on circRNAs (not conserved in multiple samples).** bsrMREs miRNA are in red. CircRNAs with multiple binding sites for the same miRNA are in blue. Some miRNAs have multiple targets.

| miRNA | circRNA | Lj gene ID | Product | Action |
|---|---|---|---|---|
| **Rhiz specific** | | | | |
| **miR11078g-3p** | LjG1.1_chr1:111879924-111880313 | LotjaGi1g1v0579100 | Protein DETOXIFICATION; AT3G08040.1 MATE efflux family protein | Cleavage |
| | LjG1.1_chr1:116785589-116787087 | LotjaGi1re1v10959800 | repeat | Cleavage |
| **miR11177-5p** | LjG1.1_chr2:94271457-94271594 | LotjaGi2g1v0449000 | Exportin-1; TAIR: AT5G17020.1 exportin 1A | Cleavage |
| | LjG1.1_chr4:25449378-25450306 | LotjaGi4g1v0153200 | Uridine kinase; AT1G26190.1 Uridine kinase family; | Cleavage |
| **miR11090e-5p** | LjG1.1_chr5:53077198-53078250 | LotjaGi5g1v0224200 | Respiratory burst oxidase-like protein; tair=AT1G09090.2 respiratory burst oxidase-like protein | Cleavage |
| **miR11100-3p** | LjG1.1_chr6:62183070-62183925 | LotjaGi6g1v0304500 | RING/U-box superfamily protein; tair=AT3G06330.1 Probable E3 ubiquitin ligase SUD1 | Translation |
| **miR11108f-5p** | LjG1.1_chr1:75073541-75073799 | LotjaGi1g1v0309100 | Adenine/guanine permease;interpro_id=IPR006043 (Xanthine/uracil/vitamin C permease); tair=AT3G10960.1 AZA-guanine resistant1 | Cleavage |

**Table 4.** (continued).

| LN specific | | | | |
|---|---|---|---|---|
| **miR11080-3p** | LjG1.1_chr4:78234312-78234386 | LotjaGi4g1v0406100 | RNA-binding protein 39; TAIR: AT5G09880.1 Splicing factor, CC1-like protein | Cleavage |
| **miR7527** | LjG1.1_chr5:43404324-43406752 | LotjaGi5re1v3656200 | LTR/Copia | Cleavage |
| **miR7518** | LjG1.1_chr6:27443272-27443422 | LotjaGi6g1v0114800 | RING finger protein; TAIR: AT3G47160.1 RING/U-box superfamily protein | Cleavage |
| **miR11067-3p** | LjG1.1_chr4:69973326-69973653 | LotjaGi4g1v0321600 | RRP12-like protein; TAIR: AT2G34357.1 ARM repeat superfamily protein; | Cleavage |
| | LjG1.1_chr5:27223731-27223890 | LotjaGi5re1v2449400 | LTR | Cleavage |
| | LjG1.1_chr4:1863649-1863779 | LotjaGi4g1v0014800 | Peroxidase; TAIR: AT4G21960.1 Peroxidase superfamily protein | Translation |
| | LjG1.1_chr5:66454616-66454783 | LotjaGi5g1v0348700 | phospholipase-like protein (PEARLI 4) family protein; TAIR: AT2G16900.1 phospholipase-like protein (PEARLI 4) | Cleavage |
| **miR171a** | LjG1.1_chr6:48404847-48405996 | LotjaGi6g1v0188800 | Kinase family protein; tair=AT5G10270.1 cyclin-dependent kinase C; Cyclin-dependent kinase C-2 | Cleavage |

**Table 4.** (continued).

| | | | | |
|---|---|---|---|---|
| **miR11072j-3p** | LjG1.1_chr1:11810436-11810606 | LotjaGi1g1v0081900 | Triose-phosphate transporter family protein; TAIR: AT3G11320.1 Nucleotide-sugar transporter | Cleavage |
| **HN specific** | | | | |
| **miR171b** | LjG1.1_chr5:41432170-41432368 | LotjaGi5g1v0161400 | ATP-dependent DNA helicase; TAIR: AT4G25120.1 P-loop containing nucleoside triphosphate hydrolases superfamily protein; | Translation |
| **miR11128-3p** | LjG1.1_chr1:120556978-120557118 | LotjaGi1g1v0647900 | Cationic amino acid transporter, putative; TAIR: AT3G03720.1 cationic amino acid transporter 4; | Cleavage |
| **miR11169-5p** | LjG1.1_chr1:5444315-5448545 | LotjaGi1g1v0039100_LC | 26S proteasome non-ATPase regulatory subunit 7 family protein; tair=AT5G05780.1 RP non-ATPase subunit 8A; | Cleavage |
| | LjG1.1_chr1:126281457-126284606 | LotjaGi1g1v0694900_LC | Major facilitator superfamily protein; tair=AT1G50870.1 F-box and associated interaction domains-containing protein; best_ | Translation |
| **miR11072i-3p** | LjG1.1_chr3:77156857-77158331 | LotjaGi3g1v0378700 | Short-chain dehydrogenase/reductase family protein; _tair=AT3G61220.1 NAD(P)-binding Rossmann-fold superfamily | Cleavage |
